# Binucleated cell formation and oncogene expression after particulate matter exposure is preceded by microtubule disruption, dysregulated cell cycle, prolonged mitosis, and septin binding

**DOI:** 10.1101/2024.01.07.574515

**Authors:** Rok Podlipec, Aleksandar Sebastijanovič, Hilary Cassidy, Lianyong Han, Pernille Danielsen, Petra Čotar, Hana Kokot, Benjamin Koroševič Koser, Ana Vencelj, Luka Pirker, Polona Umek, Gregor Hlawacek, Rene Heller, David Matallanas, Primož Pelicon, Tobias Stoeger, Iztok Urbančič, Ulla Vogel, Tilen Koklič, Janez Štrancar

**Affiliations:** Helmholtz-Zentrum Dresden-Rossendorf e.V., Ion Beam Center, Bautzner Landstrasse 400, 01328 Dresden, Germany; Jožef Stefan Institute, Condensed Matter Physics, Jamova cesta 39, 1000 Ljubljana, Slovenia; Infinite d.o.o., Zagrebška cesta 20, 2000 Maribor, Slovenia; Systems Biology Ireland, School of Medicine, University College Dublin, Belfield, Dublin 4, Ireland; Institute of Lung Health and Immunity (LHI), Comprehensive Pneumology Center, Helmholtz Zentrum München, German Research Center for Environmental Health, Ingolstädter Landstraße 1, 85764 Neuherberg, Germany; Member of the German Center of Lung Research (DZL), 81377 Munich, Germany; National Research Centre for the Working Environment, Copenhagen DK-2100, Denmark; J. Heyrovský Institute of Physical Chemistry, v.v.i., Department of Electrochemical Materials, Dolejškova 3, 182 23 Prague 8, Czech Republic; Jožef Stefan Institute, Low and Medium Energy Physics, Jamova cesta 39, 1000 Ljubljana, Slovenia

**Keywords:** TiO_2_ nanoparticles, MWCNTs, molecular initiating event, key event, mode of action, microtubule structural change, cytokinesis failure, aneuploidy, nuclear deformation, time-lapse fluorescence microscopy, helium ion microscopy, transcriptomics, proteomics

## Abstract

Several biopersistent high aspect ratio nanomaterials show pronounced pathogenic effects, from chronic lung inflammation and fibrosis to cancer. For example, asbestos fibers are classified as carcinogens, whereas the carcinogenicity of highly inflammatory multi-wall carbon nanotubes (MWCNTs), and TiO_2_ nanomaterials is still being evaluated. The exact early mechanisms of their pathogenicity towards inflammation and cancer remains uncertain, but it is likely not due to genotoxic or mutagenic activity. A proposed early mode of action that might lead to the formation of cancerous cells for asbestos fibers is the formation of binucleated and multinucleated cells, resulting in genetic instability. Here, we show that two high aspect ratio nanomaterials, MWCNTs and TiO_2_ nanotubes, which both induce chronic lung inflammation, induce very different cancer-related changes *in vitro*. TiO_2_ nanotubes – but not low aspect-ratio nanocubes of the same crystalline structure – disrupt microtubule organization and prolong mitosis, as well as deform nuclear shape, induce the formation of binucleated cells, and downregulate the tumour suppressor protein p53, whereas only the MWCNTs activate the stimulator of the interferon genes (STING) pathway, a hallmark of lung cancer. The observed differences in cellular responses to different high aspect ratio materials imply the need for assessing each nanomaterial individually with a broad range of tests rather than relying solely on a single marker or pathway, or even morphological or bulk chemical properties to infer possible carcinogenic properties.

## 1. Introduction

Ultrafine particulate matter (PM) can penetrate deep into our lungs and trigger multiple chronic diseases, including inflammation, fibrosis, and cancer ^1,2^. However, engineered nanomaterials are often commercialized without complete safety evaluation^3^. As current *in vivo* testing standards are lengthy and costly, alternative methodologies for safety assessment are desperately needed, but these require knowledge of materials’ mechanisms/modes of action (ref) – e.g. as a sequence of key events leading towards adverse outcomes. The mode of action depends on several factors such as size, shape, specific surface area, surface charge, and surface chemical properties ^4–6^, but such simple metrics alone cannot accurately predict the outcomes and replace *in vivo* tests ^7^.

Nevertheless, the consensus is emerging that several high aspect ratio materials, among which crocidolite asbestos fibres may be the most studied case, have a potential to induce different adverse outcomes, including cancer ^6^ ^8^. This observation has led to the formation of the so-called “fiber paradigm”, which emphasizes the significance of length (or rather aspect ratio, i.e. length versus diameter) and bio-persistence in pathogenicity of nanoparticles ^9^. Both properties contribute to the accumulation of dose ^10^, as insoluble fibers longer than approximately 20 μm cannot be cleared by macrophages (ref). Longer fibers generally pose greater toxicity ^11^, but even shorter fibers (less than 5 μm) are not harmless ^12^. For instance, multiwall carbon nanotubes (MWCNT Mitsui 7) with a mean length of 5 ± 4 μm have been shown to cause lung cancer in rats ^13^, despite being neither genotoxic nor mutagenic ^14^.

As a possible insight into the missing early mechanism, the carcinogenic long crocidolite asbestos fibers have been shown *in vitro* to cause the formation of cells with deformed nuclei and binuclear cells ^15^. The latter result from failed cytokinesis – the final step of mitosis when the cytoplasm of a mother cell splits into two daughter cells. Binuclear cells with duplicated genetic material (i.e. tetraploid) are known as a transitory state on the route to aneuploidy ^16,17^ characteristic for cancer cells ^18^ ^19–21^. The question of whether other high aspect ratio materials induce similar early signs of genetic instability remains to be answered.

In this work, we investigated such effects of two high aspect ratio nanomaterials, possibly carcinogenic by inhallation^22^, with different dimensions: multiwall carbon nanotubes (MWCNTs, mean length and diameter of 5 μm and 70 nm, respectively), and TiO_2_ nano-tubes (0.2 μm, and 10 nm) as well as low aspect ratio TiO_2_ nano-cubes (20 nm) with the same crystalline structure as the nano-tubes. We followed the duration of cell division and irregularities in nuclear properties (number and shape) of lung epithelial cells LA-4 with live-cell microscopy. We also analysed the expression of oncogenes and tumor suppressors, as well as materials’ binding of proteins involved in mitosis and cytokinesis, particularly associated with microtubules ^23^, as abnormalities in their organization after exposure to particulate matter knowingly lead to inflammatory responses *in vitro* and *in vivo*^24–28^.

## 2. Results and discussion

For our study, we selected three materials with different, but partially overlapping physico-chemical properties: two high aspect ratio materials – MWCNTs (mean length and diameter of 5 μm and 70 nm, respectively) and anatase TiO_2_ nanotubes (0.2 μm, and 10 nm) – and low aspect ratio anatase TiO_2_ nanocubes (20 nm). Their properties are summarised in Table 1, and morphology and crystallinity ^29^ presented with high-resolution helium ion microscopy (HIM) and scanning electron microscopy (SEM) and X-ray diffractograms in Figure 1A and Figure S1. We exposed the a LA-4 lung epithelial cells (details in Supporting information) to all tested nanoparticles (NPs) at the same surface dose (*S*_NPs_:*S*_cells_), being the most important determinant of NPs toxicity in lung studies ^30^. The used dose 10:1 corresponds typically to a few µg/cm^2^ and is within the frame of relevant occupational exposure ^31^. These doses correspond to IC50 dose, determined by WST-1 assay ^29^, whereas the PI-Hoechst 33342 assay shows high cell viability ^29,32^.

**Figure 1.**
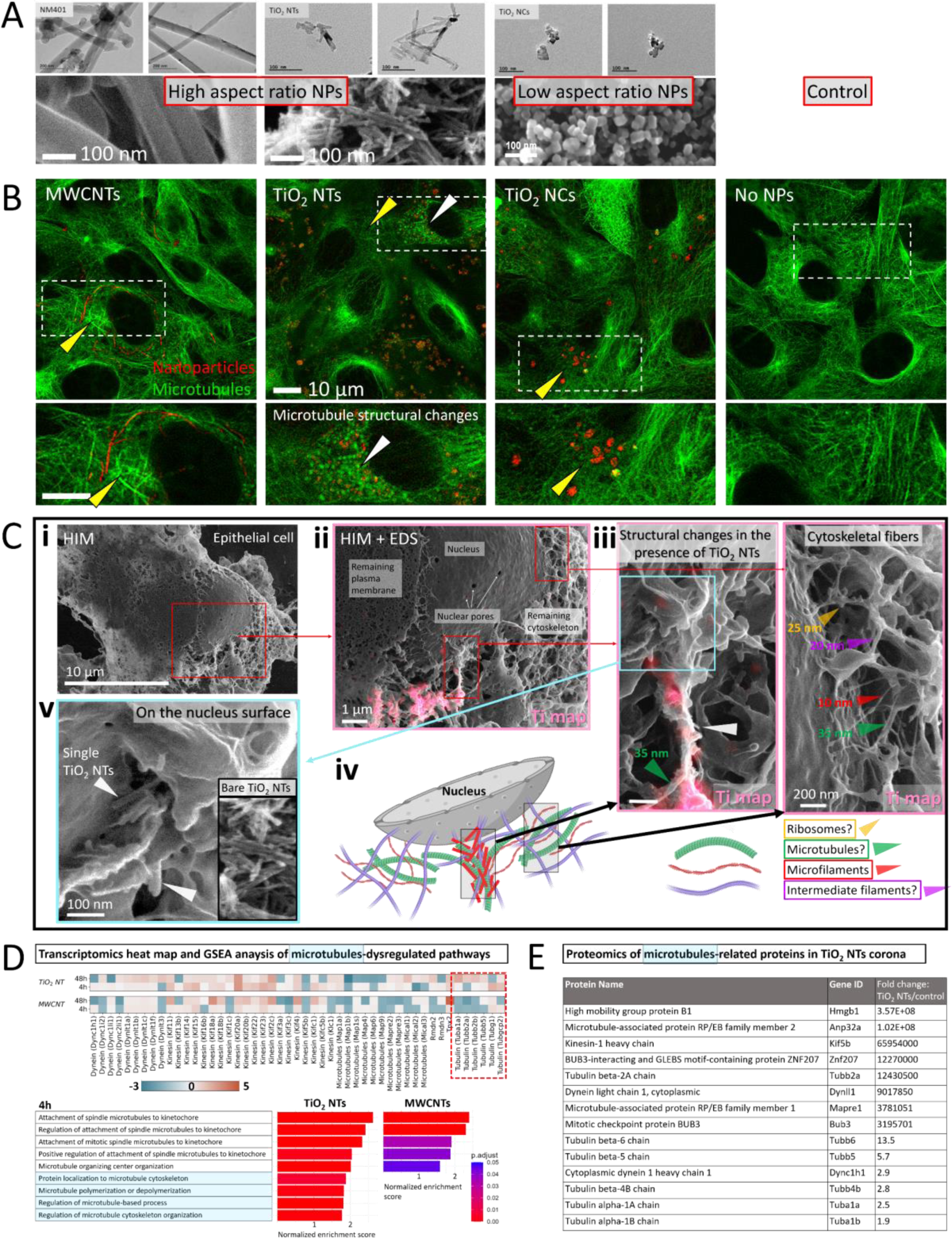
Effect of different aspect-ratio nanomaterials on microtubule structure in lung epithelial cells using confocal fluorescence microscopy and helium ion microscopy (HIM) (B-C) and supporting transcriptomics and proteomics (D-E), respectively. A) High-resolution images of tested NPs, using TEM (upper row, adapted from ^29,37^) and HIM (bottom row). B) Overlaid fluorescence (green) and backscatter (red) micrographs of microtubules and nanoparticles, with the insets showing distinct local microtubule structural changes in proximity to TiO_2_ NTs, measured 10 h after the exposure. C) i – individual lung epithelial cell exposed to TiO_2_ NTs with partially stripped plasma membrane to visualize remaining cytoskeleton measured by HIM; ii - a magnified image of the internal cellular structures showing the remaining cytoskeleton attached to the nuclear envelope, nuclear pores, and the remaining plasma membrane. Correlative HIM and Scanning Electron Microscopy-Energy Dispersive Spectroscopy (SEM-EDS) reveals the distribution of titanium (Ti) (indicated by the pink color) co-localized with the damaged cytoskeleton; iii - further magnified images reveal significant morphological changes in the cytoskeleton co-localized with TiO_2_ NTs (left, indicated by the white arrow), with microtubule-sized fiber extending out of the condensed network (indicated by the green arrow). In contrast, a preserved fiber network of all main components, intermediate filaments, microfilaments, and microtubules (indicated by color-coded arrows) is measured on the site with no NPs detected (right); iv - a schematics representing the cytoskeleton damage induced by the nanoparticles; v - a magnified ultra-high resolution image of the damaged part of the cytoskeleton bound to the nuclear envelope reveals the presence of closely packed single TiO_2_ NTs protruding out of the fiber structure (indicated by the white arrows). For more ultra-high resolution images of nuclear pores and nucleus surface with the ribosomes, please refer to Figures S8-S9 in the Supporting information. D) Transcriptomics heat map of down and up regulated expression of microtubule assembly and transport genes together with GSEA analysis of microtubules-associated upregulated pathways after cell exposure to TiO_2_ NTs and MWCNTs. The numbers represent log2 fold-change compared to the control. E) Proteomics of microtubules-associated proteins bound to TiO_2_ NTs (i.e. corona) after 2-day exposure to lung epithelial cells.

**Table 1.**
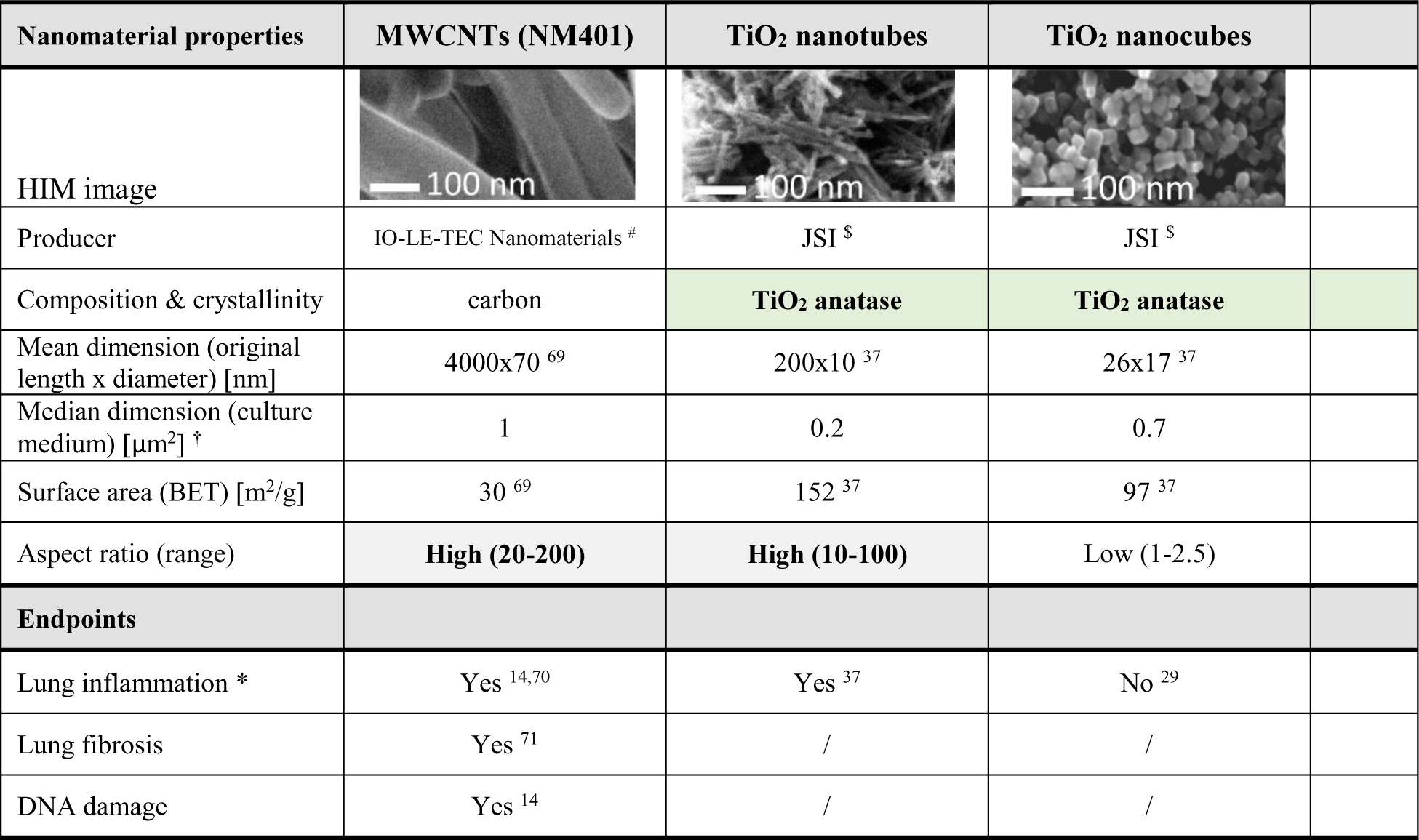
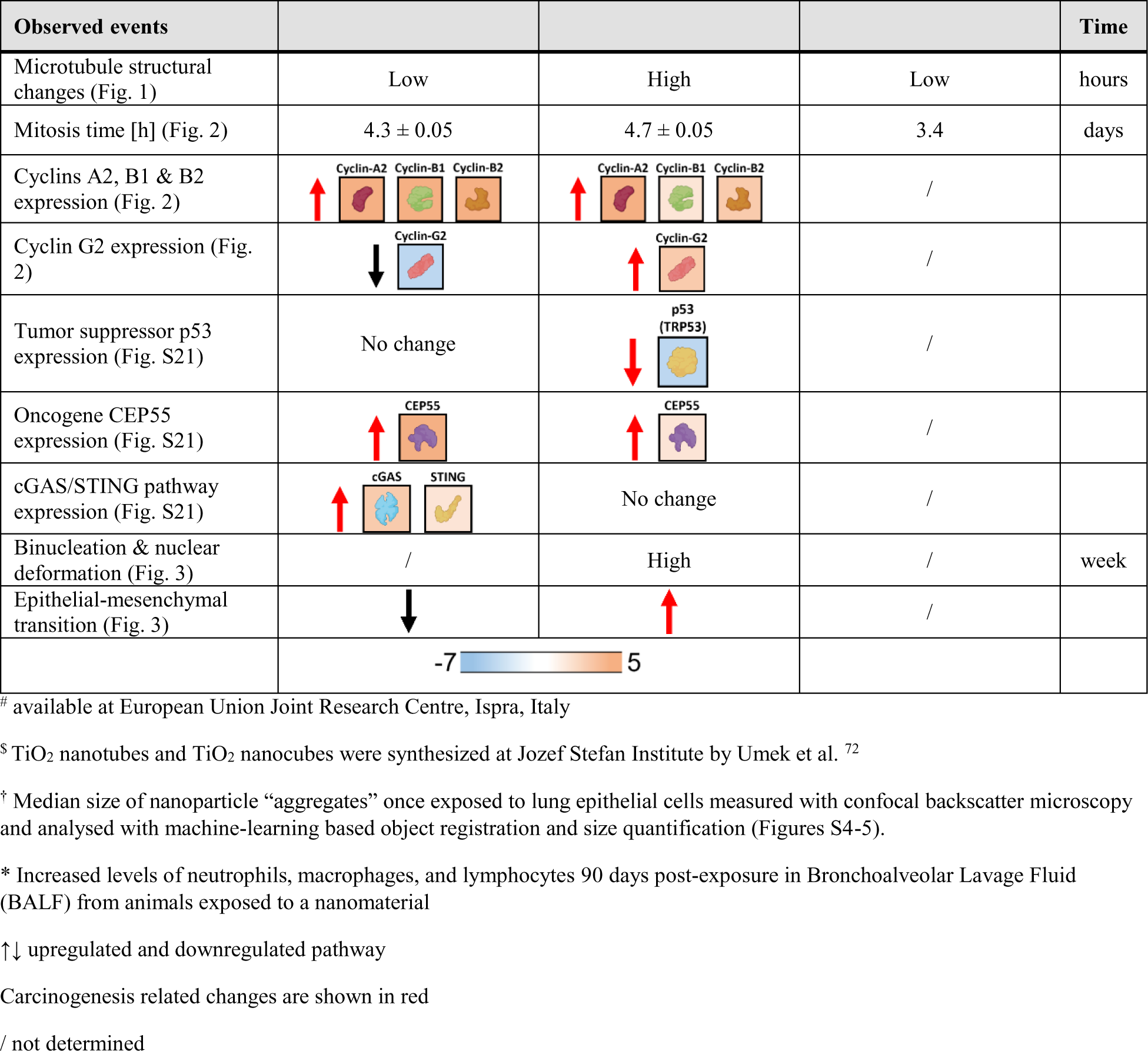
Physicochemical properties of the tested nanomaterials and their effects on lung epithelial cells in vitro after the exposure with the surface dose 10:1 (nanomaterial to cell surface). We compare two high aspect-ratio nanomaterials multi-wall carbon nanotubes (MWCNTs) and titanium oxide nanotubes (TiO_2_ NTs), as well as low aspect ratio titanium oxide nanocubes (TiO_2_ NCs) which share the same chemical and crystalline structure as TiO_2_ NTs. This comparison indicates that despite sharing some of the properties, including aspect-ratio, each nanomaterial triggers different mode of action, and suggests that comprehensive testing of each nanomaterial to establish a causal relationship between particle properties and the potential onset of carcinogenesis is needed.

### 2.1. TiO_2_ nanotubes (TiO_2_ NTs) disrupt microtubule organization

To study the impact of nanoparticles on the microtubule dynamics and organization, the key machinery for cell division, we performed time-lapse fluorescence imaging (details in Supporting Information) immediately after the exposure. Cell responses measured 10-14 hours after exposure are presented in Figure 1B with nanoparticles in red and microtubules in green. Microtubule fragmentation can be observed in proximity of the backscatter signal of TiO_2_ NTs (white arrow in the inset, see also Figures S2-S3), whereas the surrounding sites without nanoparticles mostly show intact microtubule network (yellow arrow). In addition, we could directly observe active transport of TiO_2_ NTs along the microtubule network at approx. 1 μm/min (Figure S4 and supporting Video S1), which is approximately an order of magnitude slower than the velocity of cytoskeletal motor proteins ^33^, possibly due to NPs load and disruption of protein function. n. On the other hand, we observed negligible microtubule structural changes upon interaction with the other two nanomaterials, despite high local uptake and accumulation (yellow arrows). Notably, these two materials show larger particle sizes in the cells (Figure S5), as TiO_2_ NCs seem to locally cluster inside cells (Figures S6-S7 with additional high-resolution and correlative light and ion microscopy data). MWCNTs seem to anchor and entangle with microtubule structure that could potentially disrupt cell flexural rigidity ^34^ and eventually lead to cell apoptosis ^35^. Further differences between nanoparticles are discussed in the supplementary comment #1 in Supporting information Appendix section.

For a closer look at these structures we resorted to Helium Ion Microscopy (HIM) ^36^, with sub-nanometer spatial resolution and surface sensitivity without the need for sample coating due to efficient charge compensation, preserving cellular features on the nano-scale(Figure 1C). Once identifying a cell with partially stripped plasma membrane (Figure 1C i), which occurred coincidently during the freezing of the sample, we observed distinct features of the normal cytoskeletal network (Figure 1C ii, see also Figure S8 from cells not exposed to NPs), the dimensions of which correspond to literature-reported sizes of intermediate filaments, microfilaments, microtubules (Figure 1C iii, right), as well as nuclear pores (Figure S9), and ribosomes (Figure S8). Locally to TiO_2_ NTs, identified by the pink signal from SEM-EDS elemental mapping (Figure 1C ii,iii), though, we observed a non-typical cytoskeleton network, seen as a densely packed and compressed structure indicating severe cytoskeletal damage (white arrows, Figure 1C iii, left) with the schematics of the structure presented in Figure 1C iv. By applying ultra-high-resolution imaging we also recognized single and small TiO_2_ NT aggregates through their distinct shape and dimension (200×10 nm ^37^), adhered to or impinging out of the damaged cytoskeleton in close proximity to the nuclear envelope (Figure 1C v, white arrows). ^29^ The possible mechanism of interaction is discussed in in the supplementary comment #2 in Supporting information Appendix section.

In addition, we analysed molecular transcriptomics (gene expression and Gene Set Enrichment Analysis (GSEA) ^38,39^) of LA-4 cells exposed to same doses of TiO_2_ NTs and MWCNTs for up to 48h (Figure 1D). Computational GSEA, conducted to ascertain the statistical significance of predefined gene sets presented with the heat maps revealed the TiO_2_ NTs induced clear upregulation of pathways related to microtubule (MT) assembly and polymerization, cargo transport of MT, and mitotic spindle assembly and regulation (Figure 1D). Interestingly, MWCNTs didn’t show any dysregulation of MT-based processes except the ones associated with MT spindle assembly which include upregulated expression of various microtubule motor proteins, dyneins and kinesins as shown in the heat maps measured at both time points (Figure 1D). The observed differences in the NPs impact on MT organization and its processes supports our microscopy measurements showing MWCNTs minor effect on microtubule restructuring (Figure 1B, Figure S3).

Finally, we conducted proteomics assessment of the biomaterial bound to TiO_2_ NTs particles (i.e. corona)) 48 hours following exposure to LA-4 cells measuring supernatant of exposed cells relative to the levels observed in the culture media alone. The analysis unveiled the enrichment of nanoparticles’ corona with microtubule building proteins (β-tubulin), microtubule-associated proteins relevant for microtubule polymerization/depolymerization (protein RP/EB) and cargo transport on microtubules (kinesin and dynein), and proteins involved in cell division (mitotic checkpoint protein BUB3 and kinetochore-binding protein ZNF207), crucial components in the formation of the mitotic spindle assembly ^40^. This suggests that the presence of nanoparticles with large surface area inside cells might deplete proteins relevant for the proper execution of cell division.

### 2.2. TiO_2_ nanotubes (TiO_2_ NTs) and multi-walled carbon nanotubes (MWCNTs) prolong mitosis of lung epithelial cells

The effect of TiO_2_ NTs on microtubule organization suggests that the nanotubes might also affect other cellular processes where microtubules have a key role, such as cell signaling, proliferation, motility, and division, the phenomenon already observed after exposure to other metal oxide nanoparticles ^41–43^. To quantify if the structural changes in microtubule organization are intrinsically connected to the potential changes in the cell division cycle, we conducted multiple time-lapse experiments on an expanded and stitched FoV on all studied nanomaterials and the control. We obtained a comprehensive assessment of cell division duration and cell growth for each exposed nanomaterial, as summarized in Figure 2A and Figure S10, respectively. In this context, ‘duration’ referred to the time required from cell detachment in the first stage of cell mitosis ^44^ to the central spindle disassembly in the last stage of cell division. Notably, the presented distributions of mitosis times of cells exposed to high aspect ratio nanoparticles, TiO_2_ NTs, and MWCNTs exhibited a distinct bimodal pattern, well-fitted by log-normal distribution functions (Figure 2A). The analysis performed on more than 150 cell divisions for each nanomaterial revealed some of the cells exhibit a prolonged mitosis, with the ‘prolonged mitosis component’ (designated as *t*_2_) accounting for up to 30% for MWCNTs and exceeding 1 hour for TiO_2_ NTs (depicted by orange bars and tabulated data). In contrast, the distribution of cells exposed to low aspect ratio TiO_2_ NCs did not show significant differences compared to the control. Both displayed well-fitted unimodal log-normal distributions of mitosis times (*t*_1_). The study has unveiled that the prolonged cell division cannot be unequivocally related to the microtubule disruption, meaning the mechanisms of interaction are highly intricate and fundamentally distinct for TiO_2_ NTs and MWCNTs, despite both possessing high aspect ratio shapes.

**Figure 2.**
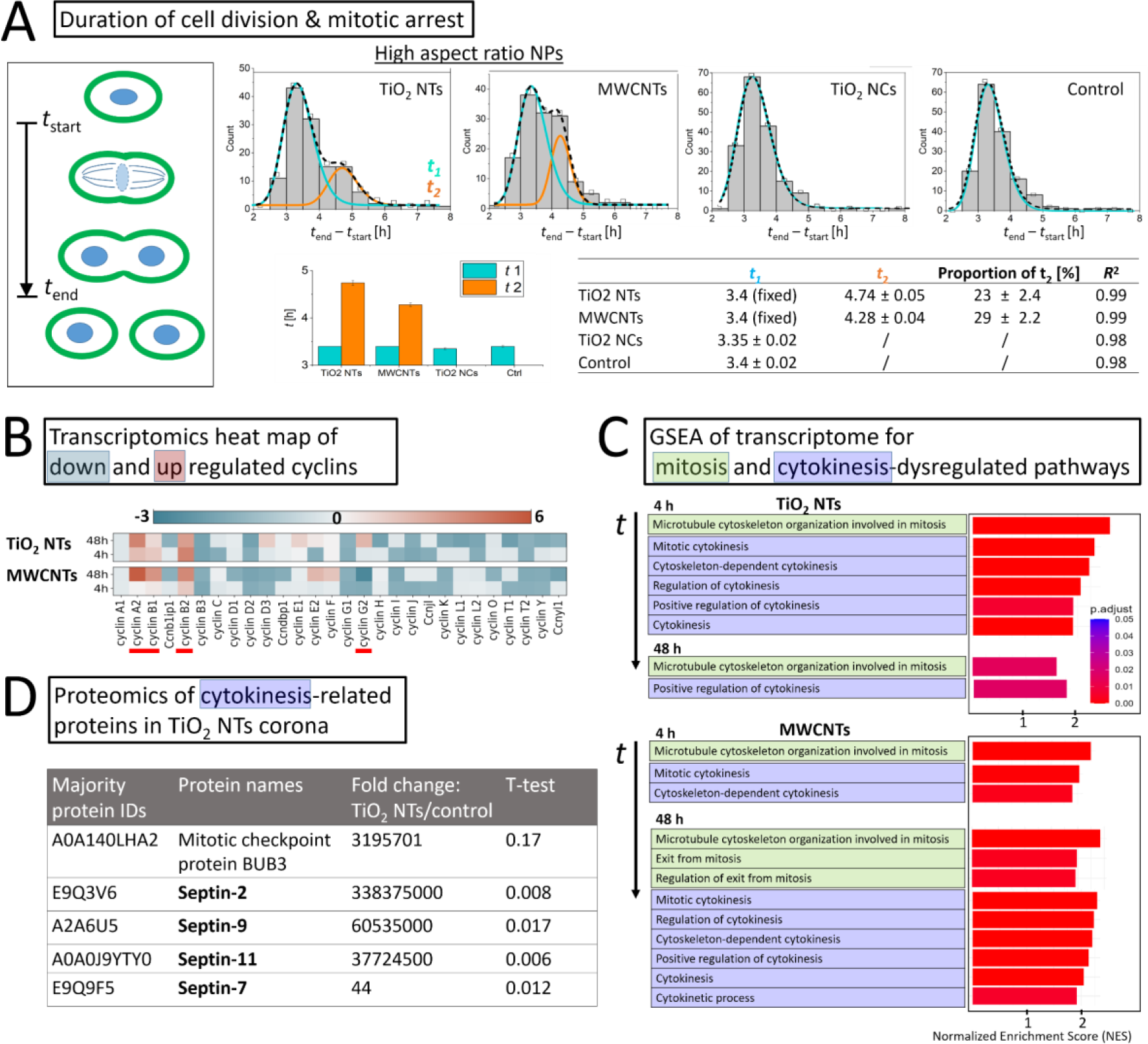
Prolonged mitosis and dysregulation of cytokinesis in lung epithelial cells after exposure to high aspect ratio TiO_2_ nanotubes (TiO_2_ NTs) and multi-walled carbon nanotubes (MWCNTs). A) The distribution of cell division times analysed from 150-250 observed events for each exposed nanomaterial exhibiting a bimodal pattern following exposure to high aspect ratio NPs, TiO_2_ NTs and MWCNTs indicating prolonged cell division (mitosis) (*t*_2_) not observed in cells exposed to low aspect ratio NPs or the control. The parameters of the log-normal component *t*_1_ were fixed to the ones measured in the control. The data was collected from 4 biological replicates. B) Transcriptomics heat maps illustrate the expression changes of cyclins, a pivotal family of regulatory proteins governing cell cycle progression, including mitosis, 4 hours and 48 hours after exposure to TiO_2_ NTs and MWCNTs at a surface dose ratio of 10:1. C) GSEA analysis reveals modifications in mitosis and cytokinesis-associated pathways after exposure to TiO_2_ NTs and MWCNTs, both of which impact the duration of mitosis. D) Proteomics analysis of cytokinesis-associated proteins within the TiO_2_ NTs corona, following a 48-hour exposure period unveiling a robust interaction with all major septins known for causing cytokinesis defects ^52^.

To further investigate the potential impact of the observed cell division arrest on cell growth, we conducted multiple 16-hour time-lapse experiments (Figure S10). The analysis of cell growth done with Cellpose algorithm ^45^, normalized to the initial cell count in the measured FoVs, did not reveal significant differences between the samples (Figure S11). It did suggest a potential slowdown after a longer period of TiO_2_ NTs exposure. We subsequently confirmed this observation through an additional multi-day experiment, presented in Figure 3B.

**Figure 3.**
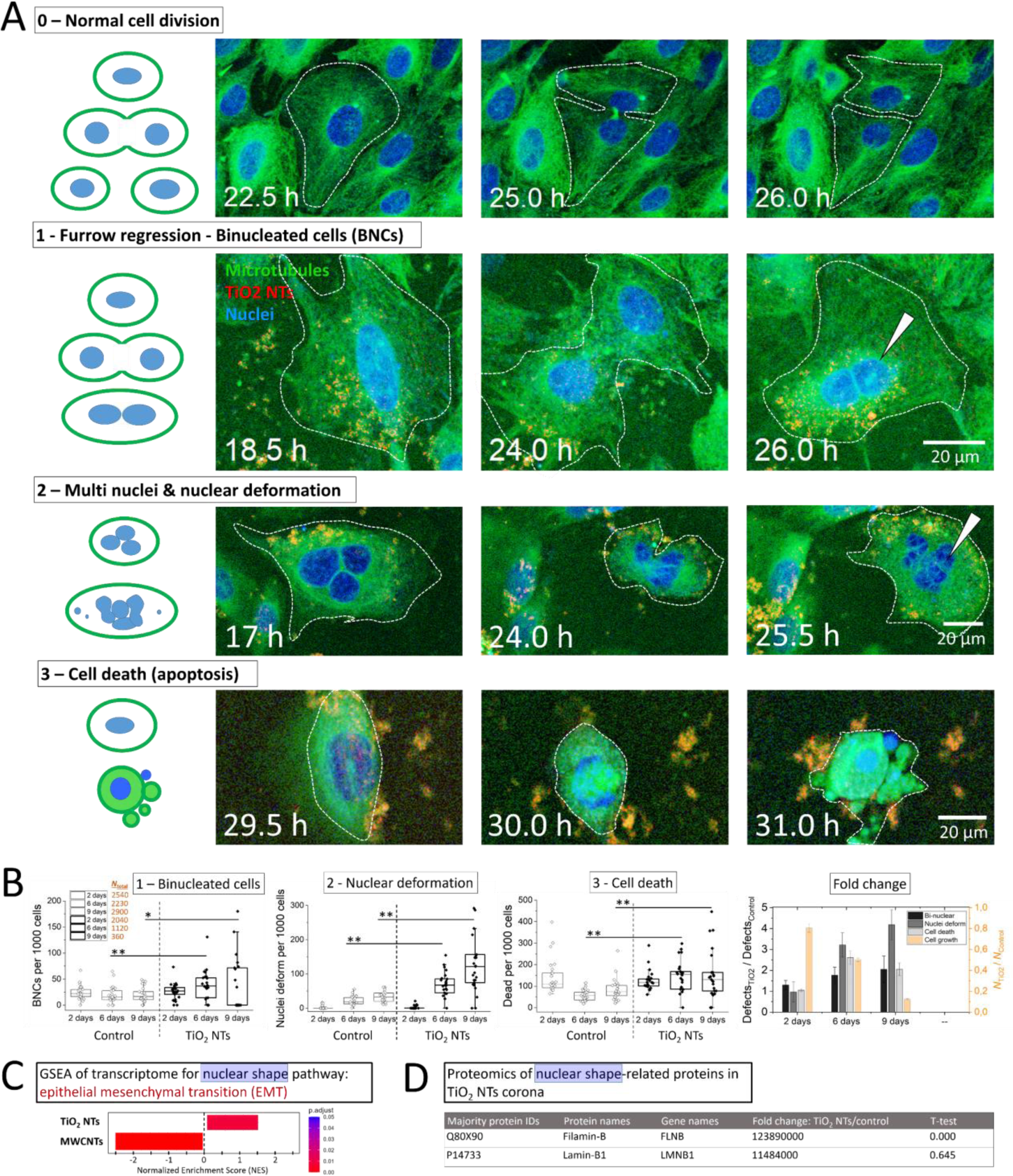
TiO_2_ nanotubes induce nuclear deformation and the formation of binucleated and multinucleated cells after 6 to 9 days epithelial cells exposure at 10:1 surface dose. A) Typical cytokinesis failures observed through furrow regression (1) (white arrows), ultimately culminating in binucleated cells (BNCs), multi(micro) nuclei formation typically accompanied with nuclear deformation (2) (white arrows) and cell apoptosis (3) that can result also from chromosomal ultrafine DNA bridge formation (Figures S19-S20). For ease of visualization, cells are outlined with dashed lines, NPs shown in red, microtubules in green and nuclei in blue. We performed the experiments using 20X objective with *NA* = 0.8. B) The frequency of the observed cytokinesis failures and cellular defects per 1000 cells on big data analysis comprising up to 3000 cells (*N*_cells_) for each sample at three TiO_2_ NTs exposure time points (left) and time-changing fold-change of the observed defects normalized to the control sample with the error bar scaled by square root of the number of measured defects for individual sample (right). We analysed 25 ROIs of large 800µm*800µm FoV using 10X objective with *NA* = 0.3. Significance of the difference is presented with one-way ANOVA test * (P < 0.05) and ** (P < 0.01). C) GSEA analysis reveals opposite regulation of nuclear shape and nuclear protein lamin related pathway, the so-called EMT transition, which is associated with cancer proliferation ^58^, following exposure to TiO_2_ NTs and MWCNTs, respectively. D) Proteomics analysis on TiO_2_ NTs corona reveals extensive binding of nuclear shape and cancer proliferation associated protein (FLNB ^59^), following a 48-hour exposure period.

For a more in-depth examination into the observed mitosis and cytokinesis arrest induced by high aspect ratio NPs, especially TiO_2_ NTs, we conducted comprehensive molecular transcriptomics (gene expression and GSEA) (Figures 2B-C) and proteomics analysis on TiO_2_ NTs coronome (Figure 2D). Briefly, transcriptomics data represented with heat maps reveals the overexpression/up-regulation of cyclins A2 and B, key regulators in the cell division cycle ^46,47^ observed in response to both TiO_2_ NTs and MWCNTs. Notably, increased expression of cyclin A2 has been previously characterized in various tumor types ^48^, including lung tumors ^49^ while dysregulation of cyclin B is known to induce centrosome abnormalities, leading to errors in chromosome segregation ^49^. The heat maps also highlight differences in gene expression between TiO_2_ NTs and MWCNTs. Specifically, we found cyclin G2 up-regulated in response to TiO_2_ NTs and down-regulated following MWCNTs exposure. Cyclin G2 is associated with centrosomes and microtubules (MT) and its overexpression has been linked to microtubule bundling, aberrant nuclei formation, and cell cycle arrest ^50^ all of which we observed in our time-lapse experiments following TiO_2_ NTs exposure. With an additional computational GSEA analysis to ascertain the statistical significance of predefined gene sets associated with mitosis and cytokinesis pathways (Figure 2C) we revealed the upregulation of distinct mitosis and cytokinesis-related pathways in response to both TiO_2_ NTs and MWCNTs. Already at the 4-hour time point post-exposure, TiO_2_ NTs were associated with the upregulation of five cytokinesis-related pathways, while later at the 48-hour mark, also MWCNTs exposure led to a significant upregulation of six similar pathways. This again suggests on different modes of action in mitosis and cytokinesis dysregulation after cell exposure to TiO_2_ NTs and MWCNTs.

To find a mechanistic explanation how TiO_2_ NTs might interfere with cytokinesis, we conducted proteomics assessment of the TiO_2_ NTs corona 48 hours following exposure of TiO_2_ NTs to LA-4 cells as described before. The analysis unveiled a significant increase in the binding of septins, a group of proteins with potential roles in cytokinesis ^51^. Among the 14 septins examined, the analysis revealed an elevation in specific septins 2, 7, 9 and 11, known to induce cytokinesis defects, ultimately leading to binucleation ^52^ (Figure 2D). The possible impact and the outcome of septins binding to TiO_2_ NTs potentially causing cytokinesis failure is discussed in the supplementary comment #3 in Supporting information Appendix section.

To test the relevance of the septins identified in the corona of TiO_2_ NTs after exposure to lung epithelial LA-4 cells (Figure 2D) as a possible mechanistic explanation for the putative carcinogenic effect of this high aspect-ratio nanomaterial, we examined the presence of septins in the corona of a known carcinogen - asbestos particles following *in vivo* exposure in rats. We have chosen asbestos as a positive control, which is known to cause aneuploidy and ultimately lung cancer ^53^. The results showed a significant increase in cytokinesis-disturbing protein septin-11 (SEPT11) ^52^ in the corona of asbestos particles 28 days after lung exposure (Figure S13). This finding confirmed our hypothesis that the depletion of septins induced by TiO_2_ NTs might contribute to cytokinesis failure, ultimately resulting in the genetic instability of exposed cells.

### 2.3. TiO_2_ nanotubes (TiO_2_ NTs) cause cytokinesis failure, formation of binucleated cells and cells with deformed nuclei

To examine whether the prolonged mitosis, cell cycle dysregulation, and septin binding by the TiO_2_ NTs leads to defects of cell division, we conducted time-lapse fluorescence measurements with live cells. Our measurements revealed the formation of two distinct cellular defects in the presence of TiO_2_ NTs, binucleated cell formation through the furrow regression (Figure 3A, 2^nd^ row, white arrow) and multinucleated cells formation typically accompanied with nuclear deformation (Figure 3A, 3^rd^ row, white arrow). We also observed cellular defects leading to cell apoptosis (Figure 3A, 4^th^ row). In case of binucleated cell formation, we observed TiO_2_ NTs localized within the cellular region of furrow ingression (Figure S14), which presumably hinders the completion of mitosis and ultimately results in the observed furrow regression. It was accompanied with the arrested mitosis and abscission delay, all being well-known indicators of mitotic slippage, which is associated with chromosome misalignment and aneuploidy ^54^. The observed nuclear morphological abnormalities could also present one of the molecular initiating events for the onset of malignancy arising from aneuploidy and/or NPs-induced changes of e.g. nuclear structural components such as nuclear lamina ^55,56^.

To assess the statistical significance of TiO_2_ NTs impact on mitotic defects, which can lead to cytokinesis failure, and potentially carcinogenicity, as extensively discussed by Lens and Medema ^16^, we conducted a multi-day exposure experiment as explained in details in Figure S15. This study covered a broad, few millimeter-sized FoV across 25 stitched ROIs (Figure S16), which provided the analysis of up to 3000 cells (*N*_cells_) for each sample, counted automatically with an Ilastik software (Figure S17) ^57^. The results demonstrated a significant increase in the occurrence of cellular defects (Figure 3B), particularly nuclear deformation and binuclearity, after 6 and 9 days of exposure to TiO_2_ NTs, with the incidence of these defects steadily rising over time. In addition, the extent of nuclear deformation as well increased with time (Figure S18). Interestingly, there was no significant difference compared to the control group after a 2-day exposure to NPs, except for a slight increase in binuclear cells, as shown in the right figure that represents the fold change in defect incidence relative to the control. Furthermore, the study revealed a notably higher frequency of dead cells following one week after the nanoparticle exposure, while there was no discernible difference from the control after a 2-day exposure. Prolonged exposure experiments also indicated a substantial impact of the nanoparticle exposure on cell growth, which decreased by more than half after one week compared to the control (depicted by the orange bars in Figure 3B).

As nuclear deformation emerged as the most prevalent cellular defect following one-week exposure to TiO_2_ NTs, we conducted GSEA analysis on transcriptomics data obtained from the same experimental setup. This analysis focused on cytoskeleton-associated pathways related to nuclear shape and the nuclear protein lamin A (Figure 3C) unveiled a significant upregulation of the Epithelial-Mesenchymal Transition (EMT) pathway, a known actor in the cancer progression ^58^. In contrast, exposure to MWCNTs led to a substantial downregulation of the EMT pathway, indicating different mechanism of action as compared to the TiO_2_ NTs. To check the plausible binding of nuclear shape-associated proteins that could additionally explain the observed nuclear defects after TiO_2_ NTs exposure, proteomics analysis on corona has revealed an extensive binding of filamin B (FLNB) (Figure 3D), whose deficiency has recently been demonstrated to play an important role in cancer proliferation ^59^.

### 2.4. MWCNTs and particularly TiO_2_ nanotubes (TiO_2_ NTs) dysregulate expression of tumor suppressor p53 and cell cycle-related oncogenes

To gain insight into a possible mode of action of carcinogenesis of the nanomaterials classified as possibly carcinogenic, we summarized nanomaterial properties and the observed nanomaterial-induced cellular physical and molecular changes in the timeline (Table 1 and supported by Figures S21-S22). It highlights different effects and possible key events (KE) of both high aspect-ratio nanomaterials and nanomaterials with identical chemical composition. Analysis of the transcriptomics data revealed different expression of oncogenes after exposure of both high aspect-ratio nanomaterials to lung epithelial cells which is presented in Table 1 and supported in the supplementary comment #4 in Supporting information Appendix section.

Two days exposure to TiO_2_ NTs showed upregulation of several oncogenes as well as downregulation of a tumor supressor protein p53 ^60^ which wasn’t the case after exposure to MWCNTs, as discusses later. In brief, the activity of tumor-suppressor p53 gene stops the formation of tumors ^61^ and is the most commonly inactivated tumor suppressor gene in human cancers ^62,63^. The downregulation of the p53 gene in cells after exposure to TiO_2_ NTs may be attributed to the observed structural change of microtubules (Figure 1). Specifically, p53 is known to be transported to the nucleus through its direct physical interaction with microtubules via its initial 25 amino acids and aided by dynein motor proteins (schematically presented in Figure S21). In the nucleus, p53 becomes active in transcriptionally activating genes related to growth arrest and apoptosis ^64^. Under diverse stresses such as DNA damage and microtubule system disintegration, p53 is typically activated ^65^. However, the disintegration of the microtubule system resulting from exposure to TiO_2_ NTs led to significant downregulation of p53 expression (Table 1). The plausible mechanism of p53-initiated events leading to the potential onset of oncogenesis after TiO_2_ NTs exposure supported by an upregulation typical oncogenic transcription factors (Table 1) is discussed in the supplementary comment #5 in Supporting information Appendix section.

Transcriptomics conducted on lung epithelial cells exposed to MWCNTs, a nanomaterial which resembles the multi-wall carbon nanotubes Mitsui-7 known to induce carcinogenicity in mice and rats ^66^, did not reveal downregulation of the tumor suppressor protein p53 (Table 1, Figures S21-S22). However, it did show upregulation of similar oncogenes and cyclins, with a few exceptions. Microscopy measurements also indicated different modes of action following cell exposure to MWCNTs, which haven’t significantly changed microtubule structure also confirmed by the minimal dysregulation of tubulin expression. On the other hand, only MWCNTs activated the stimulator of the interferon genes (STING) pathway (Table 1), a hallmark of malignant pleural mesothelioma and a critical innate imunne pathway for tumorigenesis ^67,68^. These findings underscore the complexity of these interactions and emphasize the need for better understanding of mechanisms in cancerogenesis and for comprehensive testing of each nanomaterial, using advanced microscopy and omics, to establish a causal relationship between particle properties and the potential initiation of carcinogenesis.

## 3. Conclusion

We employed a unique combination of time-lapse and super-resolution fluorescence imaging, followed by ultra-high-resolution imaging using helium ion microscopy (HIM) and SEM-EDS chemical mapping, complemented by transcriptomics proteomics and analysis, to shed light on molecular initiating events occurring after lung epithelial cells are exposed to specific nanoparticles, representing the potential mechanisms that could lead to the onset of carcinogenesis. Our findings suggest that neither nanoparticle shape, nor solely chemistry, unequivocally play the key role in inducing cytokinesis defects leading to cell deformation and genetic instability. Among the tested nanomaterials, we found TiO_2_ nanotubes (TiO_2_ NTs), but not TiO_2_ nanocubes (TiO_2_ NCs) of the same bulk crystalline structure, change microtubule organization, prolong cell division and induce mitotic arrest, commonly accompanied by cytokinesis failure via furrow regression causing the formation of binucleated cells and nuclear deformation. In addition, the proteomics showed that the TiO_2_ NTs bind high amounts of beta tubulins and septins, related to the formation of binucleated cells ^16^ and genetic instability, as observed for asbestos in rats, potentially pointing to a common interaction tendency for TiO_2_ NTs and asbestos fibers.

On the other hand, we found that similarly shaped, long aspect ratio multiwall carbon nanotubes (MWCNTs), also prolong mitosis and induce dysregulation of cell cycle and expression of oncogenes similarly as titanium oxide nanotubes TiO_2_ NTs, being in accordance with the fiber pathogenicity paradigm. However, we observed several differences in their mechanism of action, from different extent of induced microtubule structural change and regulation of microtubule-based processes to different expression of tumor suppressor protein p53, expression of the STING pathway, as well as epithelial to mesenchymal transition pathway, and nuclear shape deformation. This unambiguously indicates on high complexity of the interactions and causality of the molecular events, which requires the necessity for comprehensive testing of each nanomaterial even when possessing the similar shape or chemistry.

## Supporting information

Experimental section, Cell Culture: LA-4 murine lung cell line, µ-Slide 8 Well; Nanoparticles: Anatase TiO_2_ nanotubes (TiO_2_ NTs), anatase TiO_2_ nanocubes (TiO_2_ NCs), multi-walled carbon nanotubes (MWCNTs), BET surface area, transmission electron microscopy (TEM), powder X-ray diffraction (XRD), JRC Repository; Sample preparation for live-cell confocal and super-resolution (STED) fluorescence imaging: SiR-Tubulin, nanoparticle functionalization, Olympus IX83 with Okolab, temperature-, gas-, and humidity-controlled chamber, nanoparticles’ surface dose; Confocal laser scanning fluorescence and backscatter microscopy of microtubules and label-free nanoparticles: STED microscope and Imspector software (both Abberior), simultaneous and gated 2-channel photon detection with the avalanche photodiodes (APDs, SPCM-AQRH, Excelitas), filter settings for label-free reflectance measurements, 60x water immersion objective (Olympus UPLSAPO 60x, *NA* = 1.2); Image analysis of nanoparticles interaction with microtubules: Ilastik machine-learning-based software, training and classifying different objects, fragmented microtubules, co-localization of objects; Time-lapse (3D) multi-ROI imaging: integrated custom-built Python software, precise automation with incorporating autofocus correction, CellPose algorithm, low-magnification imaging for big data analysis of pathologies; Sample preparation for high-vacuum ultra-high-resolution microscopy: rapid cryofixation (plunge freezing), freeze drying (Coolsafe 100-9 Pro); Helium Ion Microscopy (HIM): Orion NanoFab HIM (Zeiss), sub-nanometer resolution and nm surface sensitivity, no requirement for specimen coating; Correlative scanning electron microscope energy dispersive X-ray spectrometry (SEM-EDS, Oxford instruments): Helios Nanolab 650 FIB-SEM system (FEI), CLEM Silicon holder with a silicon nitride Si_3_N_4_ membrane (PELCO); Transcriptomics sample preparation and data analysis: RNeasy Plus Mini Kit (Qiagen), Agilent 2100 Bioanalyzer, microarray analysis, WT PLUS Reagent Kit (Thermo Fisher Scientific), Mouse Clariom S arrays (Thermo Fisher Scientific), GeneChip Scanner 3000 7G, Gene Set Enrichment Analysis (GSEA), clusterProfiler package in R (version 4.0.4), Gene Ontology Biological Process (GOBP) database, normalized enrichment score (NES), log2 fold-change analysis; Proteomics sample preparation and data analysis: centrifugation and washing in acids, Thermo Scientific Q Exactive mass spectrometer, Dionex Ultimate 3000 (RSLCnano) chromatography system, a laser puller (Sutter Instruments P2000, Reprocil Pur (Dr. Maisch, Ammerbuch-Entringen, Germany) C18 reverse-phase, sample separation, Q-Exactive mass spectrometer, MaxLFQ protein quantification, Mus musculus reference proteome database (UniProt), log2 fold-change analysis, t-test significance analysis.

Supplementary Figures S1-S23: S1 - Powder X-ray diffractograms (XRDs) of TiO_2_ nanoparticles; S2 - Machine-learning based object registration and classification; S3 - The density distributions of *d*_min_; S4 - TiO_2_ NTs co-localization with microtubule network using super-resolution STED; S5 Nanoparticle size distribution during the exposure to lung epithelium LA-4 cells; S6 - TiO_2_ nanoparticles’ uptake and aggregation in lung epithelial cells; S7 - correlative high-resolution microscopy of TiO_2_ NCs internalization; S8 - Cytoskeleton structure of control sample measured with ultra-high resolution HIM; S9 - Ultra-high resolution HIM images of nuclear envelope and nuclear pores; S10 - Cell growth and typical central spindle formation in the final stage of cell division; S11 – Comparison of cell growth rate after exposure to different nanoparticles; S12 – Multiple micronuclei formation, furrow regression and microtubule disruption after cell mitotic slippage; S13 - Binding of septin-11 found in the coronome of crocidolite (asbestos) particles 28 days after lung exposure to rats; S14 – binucleated cell formation in the presence of TiO_2_ NTs; S15 Schematics of multi-day TiO_2_ NTs exposure to lung epithelial LA-4 cells; S16 - Impact of high aspect ratio TiO_2_ NTs on LA-4 cell growth on a broad field-of-view; S17 - Automated Ilastik analysis of the cell number; S18 - The time evolution of the formation and the extent of typical nuclear defects after TiO_2_ NTs exposure causing genetic instability; S19 - The formation of ultrafine DNA bridges (UFBs) during cell division; S20 - An example of cell apoptosis following chromosomal UFBs formation; S21 – The timeline of physical and molecular changes on cells after exposure to high-aspect ratio TiO_2_ NTs and MWCNTs; S22 – Graphs of dysregulated mRNA expression for cytokinesis and tumorigenesis related pathways after cell exposure to high aspect ratio TiO_2_ NTs and MWCNTs; S23 – Transcriptomics data of up- and down-regulated oncogens and oncogens found in TiO_2_ NTs corona.

## Acknowledgements

This work was supported by Helmholtz European Partnering (CROSSING project, Grant No: PIE-0007), the European Union’s Horizon 2020 research and innovation programmes (SmartNanoTox, Grant No. 686098, nanoPASS, Grant No. 101092741, HARMLESS, Grant No. 953183 and “RADIATE”, Grant No. 824096), the Slovenian research and innovation agency (ARIS) program (Experimental biophysics of complex systems, P1-0060) and ARIS Grants J7-2596, N1-0090, P1-0112, I0-0005, L7-4535 and N1-0240. The authors thank prof. Katarina Vogel Mikuš for help in sample preparation for high-vacuum measurements, Ervin Mustafic for help with image analysis and JRC Nanomaterials Repository for supplying the nanomaterials (NM-401).

## Author contributions

Conceptualization: R.P., T.K., I.U. and J.Š.; data collection: R.P. and T.K.; performed the experiments: R.P., A.S., H.K., A.V. and L.P.; data analysis and tools: R.P., A.S., H.C., L.H., P.Č., H.K., A.V., L.P., P.U., G.H., D.M., T.S. and U.V.; data visualization: R.P., A.S., L.H., P.Č., B.K.K., T.K. and I.U.; writing - original draft preparation, R.P. and T.K.; writing - review and editing: R.P., G.H., T.S., U.V., I.U., T.K. and J. Š.; funding acquisition: R.P., R.H., D.M., P.P., T.S., I.U., U.V., T.K. and J.Š.

