## Supporting Information for "Binucleated cell formation and oncogene expression after particulate matter exposure is preceded by microtubule disruption, dysregulated cell cycle, prolonged mitosis, and septin binding"

### Experimental Section

#### Cell Culture

The LA-4 murine lung epithelial cell line (ATCC CCL-196) that resemble the non-transformed alveolar type 2 epithelial cells was purchased from the American Type Culture Collection (ATCC). LA-4 cells were chosen to be a suitable model for particulate matter-induced transformation of adenoma to adenocarcinoma (cancerous cell) since they have many features of normal lung epithelial cells and are not tumorigenic but, on the other hand, have genetic characteristics of squamous cell carcinoma of the lung (L-SCC) as well as lung adenocarcinoma (LUAD). In brief, The comparison of Copy number variation (CNV) of LA-4 with specific human lung tumor types suggests that LA-4 cells demonstrate characteristic (genetic) features most consistent with non-metastatic human lung squamous cell lung carcinoma (L-SCC) [1]. Since Approximately 90% of L-SCCs are induced by mutagens, particularly toxins inhaled during smoking [2,3], it is believed that LA-4 may be considered a well-suited model for L-SCC [1]. Alveolar type 2 cells were also proposed as the cells of origin of lung adenocarcinoma (LUAD), the most common type of lung cancer [4].

The cells were cultured in 75 $cm^2$ TPP cell culture flasks and maintained at 37°C in a 5% CO2 and humidified atmosphere in F-12K medium (Gibco) supplemented with 15% Fetal bovine serum (ATCC), 1% Penicillin-Streptomycin (Sigma), and 1% non-essential amino acids (Gibco). Once the cells reached 80-90% confluency at the appropriate passage, they were seeded onto imaging holders customized for live-cell imaging (μ-Slide 8 Well, Ibidi) and correlative microscopy (Silicon Nitride Support Film, PELCO, 21509CL-10, Ted Pella).

**Nanoparticles selection, preparation and characterization**

Titanium dioxide ($TiO_2$) and multi-walled carbon nanotubes (MWCNTs), which we used in our study, have been classified by the European Commission and the International Agency for Research on Cancer (IARC) [5] as Group 2B carcinogens, meaning that they are possibly carcinogenic to humans by inhalation. Another study in genetically modified animals showed no evidence of carcinogenicity following oral administration or inhalation of $TiO_2$[6]. However, negative data from such highly sensitive genetically modified systems are considered less reliable than positive data. Studies on exposure to $TiO_2$ nanoparticles have raised real concerns for human health following inhalation or oral exposure[7,8]. In addition, ultrafine particulate matter (PM) with physical properties similar to $TiO_2$, such as large specific surface area, can penetrate deep into our lungs and has recently been shown to have carcinogenic potential in humans[9,10]. Similarly, several studies have shown the genotoxicity of certain forms of ultrafine $TiO_2$[11,12], further supporting the evidence for the carcinogenicity of $TiO_2$. Thus, the carcinogenic potential of $TiO_2$ and MWCNTs still requires further validation[13], which we address experimentally in our study.

Anatase $TiO_2$ nanotubes ($TiO_2$ NTs) and anatase $TiO_2$ nanocubes ($TiO_2$ NCs) were synthesized in-house using a method described in reference [14]. The synthesis of anatase $TiO_2$ NTs involved consecutive heating and cooling steps, transforming sodium and hydrogen titanate nanotubes into final $TiO_2$ with a diameter between 6 to 11 nm, mean length of 100-500 nm, and a BET surface area of 150 $m^2$ $g^{-1}$ [15]. In brief, 250 mg of $H_2Ti_3O_7$ nanotube sample was weighed in an alumina boat, placed into an oven (Carbolite, model CTF 12/65/550) and heated at a ramp rate of 1 °C/min to 380 °C. The sample was kept at the selected temperature for 12 hours and then cooled down to room temperature. Similarly, $TiO_2$ NC with mean dimension of 17*17 nm and a BET surface area of 100 $m^2$ $g^{-1}$ [15] were synthesized using different temperature ramps. In brief, $H_2Ti_3O_7$ nanotubes (250 mg) were suspended in 20 mL of 0.1 M solution of ethanolamine in deionized water. The prepared reaction mixture was transferred in a 30 mL glass vial and inserted into a microwave reactor (Anton Paar microwave reactor Monowave 300) and heated at 180 °C for three and a half hours under constant stirring (300 rpm). After being cooled down to room temperature the product mixture was centrifuged, washed with EtOH, dried in an oven at 100 °C overnight and then finally calcined at 280 °C for 10 h.

The morphology of $TiO_2$ was analyzed using transmission electron microscopy (TEM, Jeol 2100, 200 kV) and field emission scanning electron microscope (FE-SEM, Verios G4; Thermo Fischer). Specimens for TEM observation were prepared by dispersing a small amount of a powder sample ultrasonically in MeOH. One drop of prepared dispersion was deposited on a lacey carbon film supported by a copper grid. $TiO_2$ structural properties and crystalline phases were characterized by powder X-ray diffraction (XRD) using a D4 Endeavor, Bruker AXS diffractometer with Cu Ka radiation ($\lambda$ = 1.5406 A) and a Sol-X energy-dispersive detector. Diffractograms were measured in the 2θ angular range between 5 and 70 ° with the step size of 0.02 °/s and the collection time of 3 s (Figure S1).

Multi-walled carbon nanotubes (MWCNTs) with a diameter of 65 nm, mean length of 4000 nm, and a BET surface area of 20 $m^2$ $g^{-1}$ [16] were purchased from JRC Repository (NM-401). Nanoparticle suspensions were stored in 100X diluted bicarbonate buffer.

**Sample preparation for live-cell confocal and super-resolution (STED) fluorescence imaging**

LA-4 cells were seeded at approximately 30% confluency and $5*10^4$ cells/mL concentration in a 1.5H μ-Slide 8 Well (Ibidi) with added volume of 300 μl. The total number of cells seeded per each well was estimated between $1*10^4$ and $2*10^4$. When cells reached the desired confluency, typically in two days, they were stained with SiR-Tubulin (Spirochrome), a highly specific fluorescent probe that binds to microtubules, at a concentration of 300 nM. The staining was performed 6 hours prior to live-cell imaging. For STED fluorescence imaging of NPs co-localization with microtubules (Figure S4), $TiO_2$ NTs were labeled with STAR 580 NHS ester (Abberior) following surface functionalization with AEAPMS linker described in details in our recent paper [17]. For live-cell imaging, the samples were transferred from the cell incubator to a temperature-, gas-, and humidity-controlled chamber (Okolab, H301-MINI), which was mounted on the Olympus IX83 stage. After selecting sufficient number of the regions of interest (ROIs) (typically 4 for each sample well), time-lapse experiment was started. To introduce nanoparticles onto the cell monolayer, an appropriate volume of nanoparticle suspension ($c$ = 0.5 mg $mL^{-1}$) was carefully rinsed onto the cells, with a calculated surface dose of 10:1 ($S_{NPs}$:$S_{cells}$), assuming nanoparticles being completely dispersed. However, in realistic environments, such as cell media, complete dispersion may not always be achieved (Figure S2), resulting in a reduced effective surface dose. The suspension concentration 0.6 mg/ml was selected with a specific purpose. To achieve the desired average surface dose ($S_{NPs}$:$S_{cells}$), which is the most important determinant in the nanomaterial toxicity in lungs [18], of 1:1, 10:1 or 100:1, we calculated that 1 µl, 10 µl, and 100 µl of NPs suspension, respectively, are needed for a cell culture plate with a 1 $cm^2$ surface area. This approach allowed us to easily calculate the required suspension volume for achieving the desired surface dose in any cell culture plate and standardized our experimental protocols for these studies. For 10:1 ($S_{NPs}$:$S_{cells}$), 10 µl suspension was used, resulting in a total added surface dose of 6 µg/$cm^2$ (achieved by adding 300 µl of the final suspension to an 8-well chamber with a surface area of 1 $cm^2$). To put this into perspective, when considering the alveolar surface area of adult mice, which is approximately 80 $cm^2$ (as reported by [19]), our used dose is equivalent to exposure over a period of 45 working days, based on the 8-hour time-weighted average occupational exposure limit for $TiO_2$ established by Danish Regulations (6.0 mg/$m^3$ TiO2) [20]. The used dose is also within the frame of the relevant dose of PM2.5 (particulate matter with a diameter of 2.5 micrometers or smaller) in a polluted environment, where the surface doses can reach micrograms per square centimeter (µg/$cm^2$) in a yearly exposure. Before the exposure, the $TiO_2$ nanoparticles were gently sonicated in ultrasound bath (Bransonic ultrasonic cleaner, Branson 2510EMT) for 10 s to homogenize the dispersion.

**Confocal laser scanning fluorescence and backscatter microscopy of microtubules and label-free nanoparticles**

We performed hybrid confocal fluorescence and backscatter imaging using a customized STED microscope (Abberior) built on a fully motorized and programmable inverted Olympus IX83 microscope system. Our system featured simultaneous and gated 2-channel photon detection using two picosecond pulsed lasers, emitting at 640 nm and 518 nm wavelengths for fluorescence and backscatter imaging, respectively. The lasers provided up to 100 µW power in the sample plane, and were coupled with two high photon detection efficiency avalanche photodiodes (APDs, SPCM-AQRH, Excelitas) and a fully controlled fast FPGA board and Imspector software (Abberior). We set the pulse repetition rate to 80 MHz, with 10 µs dwell/exposure time at each image voxel, and used line scanning mode with the quad galvo scanner system (Abberior). To enable simultaneous multichannel detection, we used motorized notch filters in the excitation-detection path and band-pass filters (650–720 nm and 500–500 nm, both Semrock) installed before the detectors. Unlike complete blocking of reflected/backscattered excitation light in the fluorescence channel, the proper set of consecutive notch, dichroic, and band-pass filters enabled partial transmittance, with the total reflectance signal at the detector comparable to the fluorescence signal. We used average laser powers of 20 µW and 5 µW for fluorescence and backscatter measurements, respectively, and a 60x water immersion objective (Olympus UPLSAPO 60x, $NA$ = 1.2) for all high-resolution experiments.

**Image analysis of nanoparticles interaction with microtubules**

To quantify microtubule disruption in the presence of nanoparticles, we utilized image analysis with Ilastik, an interactive machine-learning-based software [21], presented in more detail in Figure S4. Our approach involved training and classifying different objects based on their shape and intensity, with a focus on identifying fragmented microtubules and nanoparticles. After discriminating the objects presented by color coding, red for nanoparticles and green for "fragmented" microtubules, we used their center-of-object coordinates to identify the closest pairs ($d_{min}$). To quantify spatial relation, we counted the number of co-localized objects ($d_{min} < 1\mu m$). Since microtubule structural changes can also occur independently of nanoparticle proximity, we only selected and counted objects with distances smaller than one micrometer, for which we considered to be the best representatives of the interacting nanoparticles and microtubules. To ensure statistical significance, we conducted multi-ROI analysis at 8 measuring sites for each time point.

**Time-lapse (3D) multi-ROI fluorescence imaging and analysis**

To study the dynamics of microtubule structural changes, and cell division processes divided on mitosis and cytokinesis following nanoparticle exposure, a custom-built Python software integrated with the Imspector software was used. This integration enabled precise automation of time-lapse 3D multi-ROI imaging while incorporating autofocus correction to account for any stage drift that occurred during the experiment. To ensure robustness and reliability, the cells exposed to nanoparticles and control samples were assessed in triplicate (control in duplicate) and housed in μ-Slide 8 Well chambers. To better understand the mechanisms underlying the observed effects of microtubule disruption, we measured four regions of interest (ROIs) in individual chambers at three different planes, with a 2 μm z-step at high magnification (60X), allowing for an accurate characterization of microtubule organization in both 2D and 3D. We captured 17 time points for each ROI, with a time step of 1 hour, for a total experiment duration of 16 hours. To minimize laser photo-toxicity and photo-bleaching [22], and to maintain cell function throughout the time series, we scanned each ROI fewer than 70 times.

To quantify the impact of particularly high-aspect ratio nanoparticles, on cell division processes and cell growth, we conducted additional imaging experiments to acquire sufficient statistical data for a comprehensive analysis. The data for the duration and the observed defects of cell division were collected from the time-lapse multi-ROI imaging (total of 8) using 20X objective performed in quadruplicate enabling more than 150 observed cell divisions for each sample. An example of cell growth together with a typical cell mitosis feature of cells exposed to MWCNTs and $TiO_2$ NTs is presented in Figure S12. Rate of cell growth (Figure S13) was measured within the first 16 hours of their exposure to each nanomaterial using an automated CellPose analysis for nucleus detection (https://github.com/MouseLand/cellpose) [23]. The data was fitted with the exponential curve to obtain the growth rate parameter $\tau$.

For a more detailed assessment of the impact of $TiO_2$ NTs, the nanomaterial with the most pronounced disruption of microtubules and the formation of binucleated cells, 9 days multi-ROI imaging using a 10X objective ($NA$ = 0.3) was conducted (Figures S17-18). The measurements were performed in three time points, 2 days, 6 days and 9 days of NPs exposure, each comprising 25 ROIs with the FoV 800*800 μm, yielding datasets of up to 3000 analysed cells. The number of cells was calculated using Ilastik software, which was trained to register and characterize labelled nuclei of live cells, the similar approach being used for the characterization of co-localization of NPs and the "fragmented" microtubules (Figure S5). In all experimental setups, cells were exposed to a surface dose of 10:1 ($S_{NPs}$:$S_{cells}$) prior to the measurements. Fluorescence imaging was done by labeling microtubules (SiR-Tubulin, 500 nM) and cell nuclei (Spy DNA 555, 500 nM) two hours prior to imaging.

**Sample preparation for high-vacuum ultra-high-resolution microscopy**

A 10 μL suspension of LA-4 cells at approximately 50% confluency was poured onto a UV-sterilized CLEM Silicon holder with a silicon nitride $Si_3N_4$ membrane (PELCO). After allowing sufficient time for cell adhesion on the $Si_3N_4$ membrane (typically a few hours), the sample holders were placed in μ-Slide 8-Well Chambers (Ibidi) and rinsed with cell growth media to cover the entire bottom surface of the chamber. This step was important to prevent any evaporation during the experiment. After allowing sufficient time for cell growth (typically three days), the same volume of nanoparticle suspension ($c$ = 0.5 mg $mL^{-1}$) was carefully added to the chamber to deliver the same surface dose 1:1 ($S_{NP}$:$S_{cells}$). To visualize early events of microtubule interaction with nanoparticles at high-vacuum and nm resolution, rapid cryofixation was performed using plunge freezing in liquid propane ($T$ = -185 °C) one hour after exposure, corresponding to the second time point in the time-lapse experiment. Propane was prepared in a liquid nitrogen-cooled chamber with the plunge freezer transfer system (37015, Electron Microscopy Sciences). To prevent crystalline ice phase formation, samples were rapidly cooled at a rate of $>10^4$ K/s [24]. Chemical fixation was not performed before the freezing step to preserve the integrity and morphology of cellular structures and interacting nanomaterial composites/aggregates. To provide the insight into cellular internal structures such as cytoskeleton organization and to potentially identify the internalized nanomaterial and its interaction with the structures, an excess water was removed by blotting only the sides of the sample holder with a lint-free paper, reducing the risk of contamination, for an extended period (30 s instead of typically 10 s). This allowed just enough drying to introduce local plasma membrane rupture enabling direct visualization of the internal structures after cryo fixation, while avoiding severe plasma membrane damage. Following fast plunge freezing, the samples were stored in cryo vials and immediately transferred to a cryo bank until the final drying step in a freeze dryer (Coolsafe 100-9 Pro), which was set to two days with the preset temperature ramp. Once dried, the samples were ready for high-vacuum ultra-high-resolution helium ion microscopy (HIM) and scanning electron microscopy (SEM). The sample holders were handled with PTFE-coated high-precision and ultrafine tweezers (EMS, 72919-3SATe).

**Helium Ion Microscopy (HIM)**

To achieve sub-nanometer resolution and nm surface sensitivity without requiring biological specimen coating, we utilized the Orion NanoFab HIM (Zeiss), which provides superior imaging capabilities for single nanoparticle detection [25]. The HIM incorporates a Gas Field Ion Source (GFIS) injection system that produces helium ions with a low energy spread (< 1 eV). By carefully adjusting the electrostatic lenses, quadrupoles, and octapoles, the HIM achieves a large depth-of-field (DoF) and lateral resolution as low as 0.5 nm, which surpasses the capabilities of SEM. Prior to imaging, freeze-dried samples were attached to carbon tape and mounted onto standard pins in a multi-pin specimen mount. Secondary electrons (SE1) emitted from the few nanometer surface layer were detected using an ion energy of 30 keV, ion current of 200 fA, and chamber vacuum of $3x10^{-7}$ hPa. The acquired images had a field-of-view (FoV) that varied from a lower magnification of 30*30 $\mu m^2$ to an extreme high magnification of 0.4*0.4 $\mu m^2$ with a minimal pixel step < 1 nm.

**Correlative SEM-EDS**

The Helios Nanolab 650 FIB-SEM system (FEI) was used for correlative imaging and elemental analysis. To measure the same cell structures as with HIM, CLEM holders with $Si_3N_4$ membranes were used as fiducial markers. Prior to imaging and spectroscopy, samples were coated with a 5-10 nm carbon layer to prevent charging of highly insulating biological specimens. Immersion mode settings were used for imaging, with an electron acceleration voltage of 5-10 kV, electron current of 200 pA, and secondary electron (SE) detection using the in-lens detector (TLD-SE) to maximize surface sensitivity. Energy-dispersive X-ray spectroscopy (EDS) was performed using the X-Max SDD detector (Oxford Instruments) to confirm and characterize the elemental distribution of $TiO_2$ NTs in relation to the surrounding cell structures at the observed sites. The electron current was set to 400 pA, and electron acceleration voltages were set to 10 kV to detect the Ti Kα line.

**Transcriptomics sample preparation and data analysis**

Cells were grown in 6-well plates and exposed to $TiO_2$ NTs and MWCNTs for 4 and 48 hours. After the exposure period, growth medium was removed and the cells were frozen at -70°C for further analysis as previously described [16]. Total RNA was extracted using the RNeasy Plus Mini Kit from Qiagen. RNA quality was assessed using the Agilent 2100 Bioanalyzer, and RNA with a RIN (RNA Integrity Number) > 7 was selected for microarray analysis. For the analysis, a total of 120 ng of RNA was amplified using the WT PLUS Reagent Kit from Thermo Fisher Scientific Inc. (Waltham, USA). The amplified cDNA was then hybridized onto Mouse Clariom S arrays (Thermo Fisher Scientific). Staining and scanning were conducted following the manufacturer's instructions using a GeneChip Scanner 3000 7G.

The results were represented with the heat maps showing only up- and down- regulated genes for more than two times compared to non-exposed cells. The numbers show a log2 fold-change compared to the control. The expressed genes were used for Gene Set Enrichment Analysis (GSEA) to characterize the up- and down- regulated pathways related to microtubule fragmentation, cytoskeleton modification and cytokinesis failure after $TiO_2$ NTs and MWCNTs exposure. GSEA was performed with clusterProfiler package in R (version 4.0.4) [26,27], comparing the Gene Ontology Biological Process (GOBP) and Kyoto Encyclopedia of Genes and Genomes (KEGG) database. Bar plot was used to visualize enriched terms according to the use of ggplot2 package. The width of each bar represents normalized enrichment score (NES) while color of bar plot, the adjusted *p* value. All the enriched terms are predicted significantly regulated by nanomaterial treatments ($p$ adjust $< 0.05$). Array data of $TiO_2$ part has been submitted to the GEO database at NCBI.

**Proteomics sample preparation and data analysis**

Sample preparation for the identification of nanoparticle coronas was done as previously described [28]. In summary, samples were centrifuged at 2400 relative centrifugal force (RCF) for 15 minutes in a refrigerated centrifuge. The resulting pellets contained the nanoparticle-corona complexes. After removing the supernatant, they were resuspended in 1 ml of PBS. This washing process was repeated three times to remove contaminant proteins. Clean pellets were resuspended in a buffer containing 8M urea and 100mM Tris at pH 7.5. Samples were then incubated and mixed in a thermomixer for 10 minutes at 50°C. Proteins were alkylated using 0.5 M iodoacetamide (IAA) to a final concentration of 0.05 M IAA. After incubation at room temperature in the dark for 30 minutes, the samples were diluted in a 40% acetonitrile (ACN) solution to achieve a final concentration of 2 mM urea and 10% ACN. Tryptic digestion was carried out overnight at 37°C by incubating the samples with 8 µl of sequence-grade trypsin diluted in 1 mM HCl to a concentration of 12.5 ng/µl. The next day, 100 µl of trifluoroacetic acid (TFA) was added to achieve a final concentration of 5%, and the peptides were collected by centrifugation at 24,000 RCF for 30 minutes. The supernatant was collected into a clean tube, and the pellet was resuspended in three volumes of a 50% ACN/0.5% TFA solution, followed by two additional centrifugation steps. The sample was then concentrated to a volume of 5 µl using a CentriVap concentrator, and 100 µl of 0.5% TFA was added. Filter-aided sample preparation, as previously reported [29], was performed using C18 stage tips for desalting, following the procedure described in [30]. Eluates were evaporated using a CentriVap concentrator, and the dry pellets were resuspended in 15 µl of 0.1% TFA and used for mass spectrometry analysis.

Each sample was analyzed in triplicate using a Thermo Scientific Q Exactive mass spectrometer. The settings employed were consistent with those previously reported [28]. The mass spectrometer was coupled to a Dionex Ultimate 3000 (RSLCnano) chromatography system, with the samples loaded onto a fused silica emitter with a 75 µm inner diameter. Sample loading was performed using a laser puller (Sutter Instruments P2000, Novato, CA, USA). The samples were packed with Reprocil Pur (Dr. Maisch, Ammerbuch-Entringen, Germany) C18 reverse-phase media, with a particle size of 1.9 µm and a column length of 12 cm. Subsequently, the samples were separated using an acetonitrile gradient that increased over 60 minutes, all at a flow rate of 250 nl/min, leading directly into the Q-Exactive mass spectrometer. The mass spectrometer operated in positive ion mode, with a capillary temperature of 320°C (S-38) and a potential of 2300 V applied to the frit. Data acquisition was performed in automatic data-dependent switching mode, with an initial high-resolution MS scan (300-1600 m/z) using the Q Exactive to select the 12 most intense ions before proceeding to MS/MS analysis employing high-energy collision dissociation (HCD). The MS scan operated at a resolution of 70,000.

Proteins were identified and quantified by MaxLFQ [31] by searching with the MaxQuant version 1.5 against the Mus musculus reference proteome database (UniProt). Modifications included C carbamylation (fixed) and M oxidation (variable). Excel was employed to further analyze the MaxQuant data, using fold change cut-off values of 2 folds and tests to determine significance using t-test as described before [32].

### Appendix

#### Supplementary comment #1

The reason for different cellular response to $TiO_2$ NCs compared to $TiO_2$ NTs could be attributed to the different modes of action originating in different aspect ratios, sizes, surface defects/charge [33], where $TiO_2$ NCs are predominantly internalized by the cells, forming local aggregates that reduce the available surface area for interactions (Figures S6-S7). In contrast, $TiO_2$ NTs were partially aggregated and quarantined on the cellular surface (indicated with asterisk) and partially internalized, forming on average smaller aggregates that maintained a higher surface area available for interactions. In combination with their distinct rod-like shape [34], this may cause faster rate of internalization via passive diffusion/uptake [35]. The uptake can start already seconds after the exposure as shown in our recent experiment on air-thin layer of water-epithelium interface [16]. More effective internalization in combination with the available surface area, in turn, increases the likelihood of physical interactions with the cellular machinery. To translate this visual observation into a quantifiable metric of particle size distribution once exposed to cells, we performed advanced automatic image analysis across 8 regions of interest (ROIs), from duplicate time series comprising approximately 30-40 cells in total for each sample and time point at an approximate local confluency of 80-90%. The analysis indeed revealed 3 and 6 times smaller median size of $TiO_2$ NTs compared to $TiO_2$ NCs and MWCNTs, respectively (Figure S4). The analysis also revealed that the highest microtubule fragmentation corresponds to the exposure of particularly $TiO_2$ NTs. The analysis of the microtubule fragmentation using Ilastik software [21] and the results of co-localization between fragmented microtubules and nanoparticles are presented in more detail in the Supporting information (Figures S2-S3).

**Supplementary comment #2**

These measurements provide direct evidence of single- or small $TiO_2$ nanotube aggregates-induced cytoskeleton and microtubule damage through their strong adhesion and capability of local bridging and compressing fiber structures seen as early as one hour after the exposure. Based on high-resolution microscopy and molecular transcriptomics results (next paragraph), we formulated a possible hypothesis concerning the mechanism of interaction. Microtubule exterior polymers/monomers are characterized by a net negative charge [36], while microtubule ends proximal to centrosomes are likely to exhibit positive charge [37]. Given the negative charge of $TiO_2$ NTs [16], it is conceivable that the interaction may be more localized at the microtubule ends. Consequently, this localized interaction could potentially impact the microtubules' function in association with the centrosomes—a vital organelle in the intricate process of cytoskeleton organization, microtubule disassembly/assembly, and cell division, including chromosome movement.

**Supplementary comment #3**

The depletion of septin-7, the 17$^{th}$ most abundant protein within the $TiO_2$ nanotubes' coronome, and of septin-2 have been associated with defects in chromosome alignment at the metaphase stage [38]. It appears that the $TiO_2$ NTs exhibit the capability to deplete these septins responsible for cytokinesis. Depletion of septins could be one of the key mechanisms to even cause cytokinesis failure, resulting in the furrow regression (Figure 3A) and the formation of multiple nuclei (Figure S12).

**Supplementary comment #4**

The transcriptomics analysis identified up-regulation of distinct oncogenes, notably Rab genes, recognized for their role in regulating survival pathways in cancer [39]. While up-regulation increased with the time of $TiO_2$ NTs exposure, it significantly decreased with time of MWCNTs exposure to lung epithelial cells, again indicating different mechanisms of action (Figure S23A). Notably, we also identified oncogenesis-associated Rab proteins on the corona of $TiO_2$ NTs after 4 and 48 hours of exposure (Figure S23B).

**Supplementary comment #5**

Focusing on the mechanisms, the p53 performs the crucial function of binding to DNA. It then initiates the production of another protein known as cyclin-dependent kinase inhibitor 1 (CDKN1), often referred to as p21. This p21 protein interacts with another protein responsible for promoting cell division, cyclin-dependent kinase 2 (cdk2), essentially acting as a 'stop signal' for cell division [40]. P53 downregulation leads to the upregulation of several transcription factors, including Forkhead box M1 (FOXM1), and genes associated with cell cycle progression and oncogenesis, such as polo-like kinase 1 (PLK1), centrosomal protein 55 (CEP55), Cyclin B1, and others [41], which is also observed in our study (Figures S21-S22). PLK1 is frequently identified as an overexpressed oncogene across a broad range of human tumors. This overexpression leads to abnormal chromosome segregation and interferes with cytokinesis, the final phase of cell division responsible for equally dividing the cytoplasm between two daughter cells. This disruption generates polyploid cells [42]. CEP55 is a protein associated with the centrosome and midbody, serving as a critical regulator of cytokinesis [43]. Studies have shown that overexpression of CEP55 correlates with tumor aggressiveness and metastasis, underscoring its importance in cancer progression [41]. FOXM1 is an oncogenic transcription factor with increased expression in most cancer types, while it is typically absent or present at very low levels in most normal, non-regenerating adult tissues [44]. Cyclin B1 plays a pivotal role in facilitating cell cycle progression to enable mitosis and is known to be overexpressed in various cancer types, often associated with poor prognosis [45].

### Supporting Figures

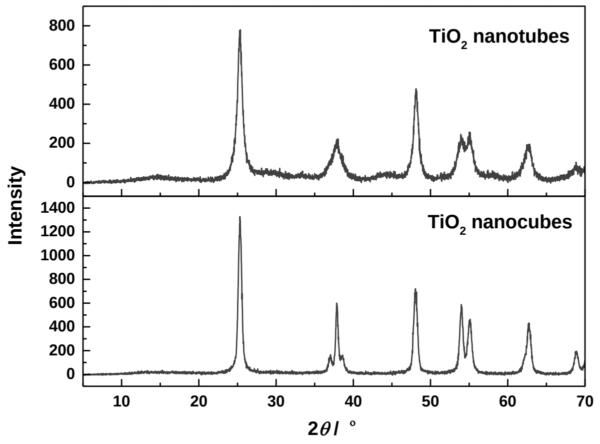


**Figure S1**. Powder X-ray diffractograms (XRDs) of $TiO_2$ nanotubes (top) and nanocubes (bottom). All diffraction peaks in both diffractograms are assigned to anatase (ICCD Card No. 86-1157).

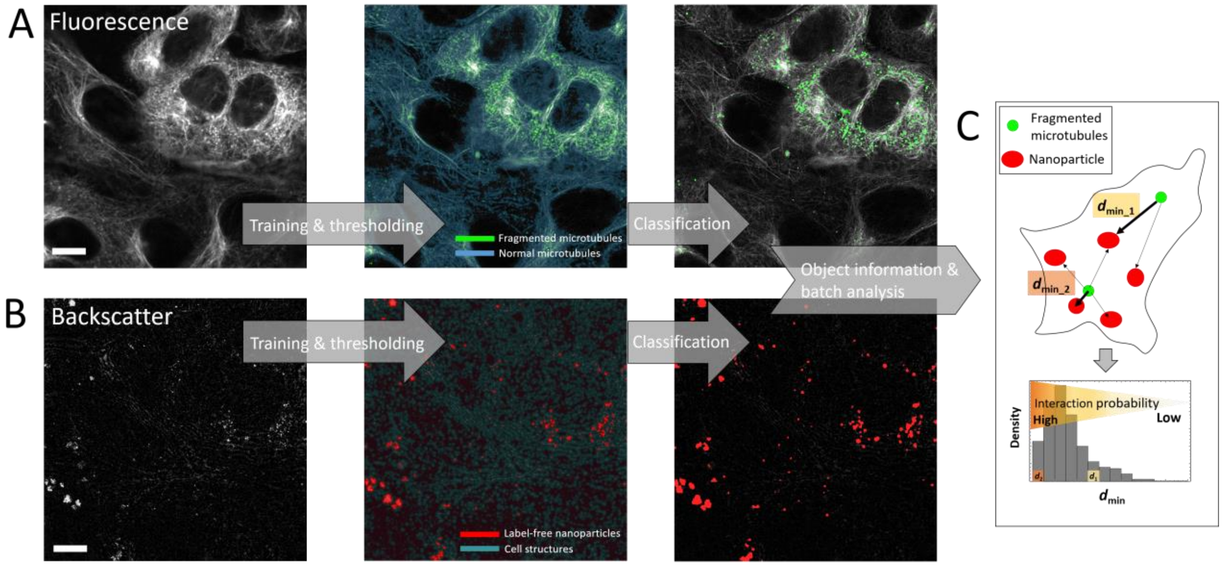


**Figure S2.** Detailed steps of machine-learning based object registration and classification (Ilastik software) for characterization of co-localization between fragmented microtubules and nanoparticles. Analysis of the fluorescence signal of microtubules (A) and backscatter signal of nanoparticles (B) with object training, thresholding, classification and final analysis (C). Scale bar is 10 μm.

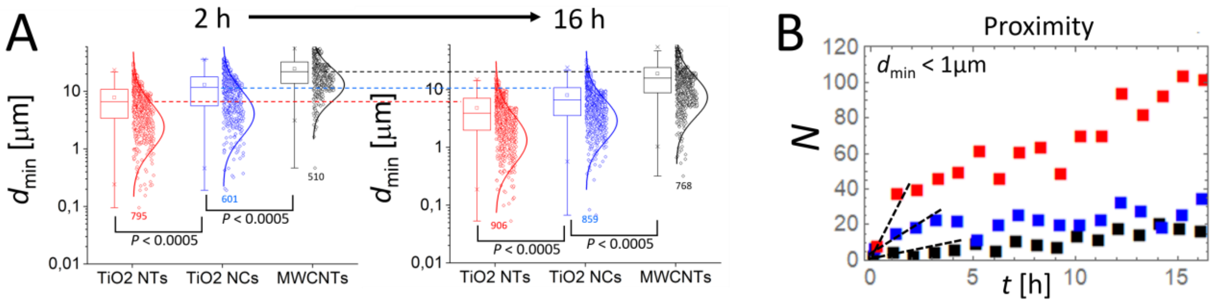


**Figure S3.** The distributions with the statistical analysis (one-way ANOVA) of $d_{min}$ obtained from multi ROI imaging 2 h and 16 h after NPs exposure (A), and the time-changing rate of the number of fragmented microtubules in proximity of NPs ($d_{min}$ < 1μm) (B). The analysis at two distinct time points after the exposure reveals significant difference in $d_{min}$ distribution for each nanomaterial, while all demonstrating a shift to lower $d_{min}$ with time (dashed lines), indicating a decrease in the average distance between fragmented microtubules and the nanoparticles. Notably, the smallest mean $d_{min}$ dropping below 5 μm, was identified with TiO2 NTs, indicating the greatest impact on microtubule dynamics and structure among tested nanoparticles (NPs). We also observed the highest Increase in the number of fragmented microtubules in proximity of TiO2 NTs during the initial time points after exposure as shown in (B).

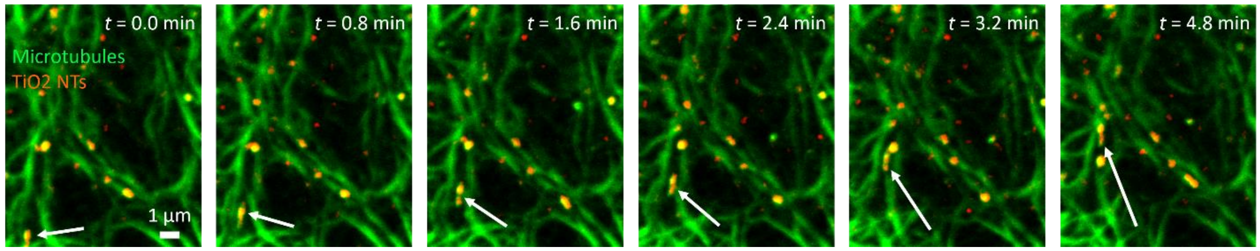


**Figure S4.** TiO2 NTs co-localization and interaction with microtubule network measured with STED super-resolution imaging (see the moving particle highlighted with white arrow). Imaging was performed with a pixel step size of 30 nm, employing a 775 nm STED laser with a power output of 100 mW.

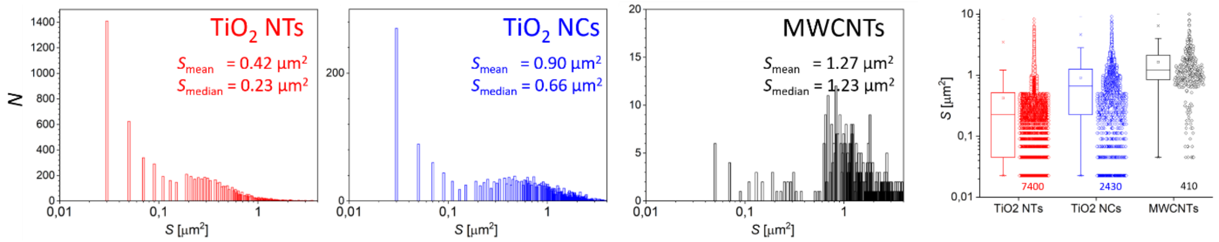


**Figure S5**. Nanoparticle size distribution with mean and median values during the exposure to lung epithelium LA-4 cells measured with label-free confocal laser backscatter microscopy using 60X magnification and *NA* = 1.2. The imaging spatial resolution enabled size characterization down to 0.02 μm$^2$.

The analysis revealed a huge difference in size distribution above 0.023 μm² among the various measured nanomaterials, as visually presented on the right. Size distributions were analyzed from 8 regions of interest (ROIs) and a first few time points after NPs exposure.

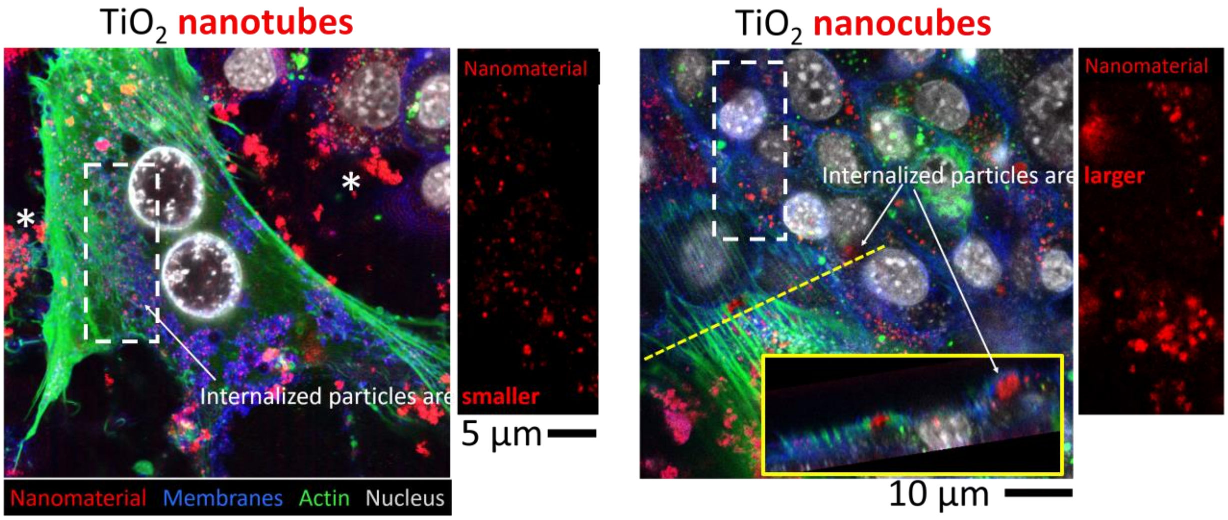


**Figure S6.** Different shaped TiO2 nanoparticles' uptake and aggregation in lung epithelial cells measured 3h after exposure measured by confocal fluorescence microscopy (CFM) using a 60X magnification objective with *NA*=1.2. Cellular structures were fluorescently stained with 1 μM CellMask Orange (all membranes), 500 nM SiR Actin (actin) and 1 μM Hoechst (nucleus) while NPs were detected by label-free confocal backscatter microscopy.

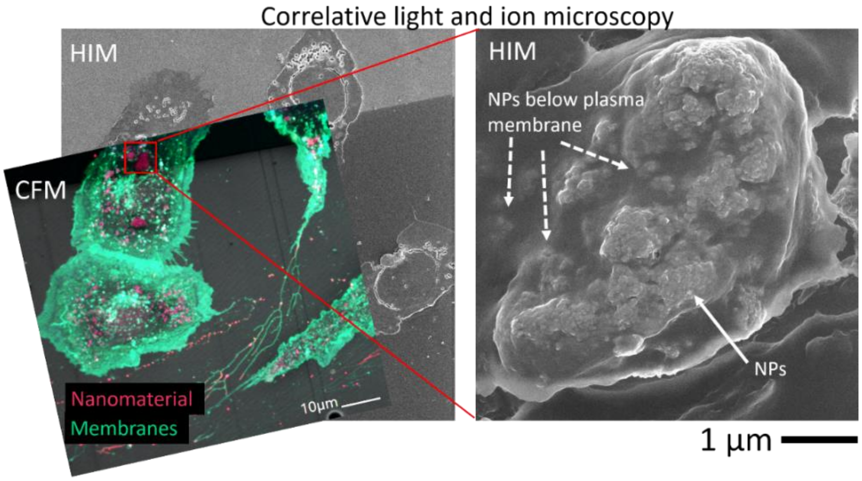


**Figure S7.** Correlative live-cell confocal fluorescence microscopy (CFM) and high-vacuum helium ion microscopy (HIM) revealing detailed morphology and extensive aggregation of $TiO_2$ NCs inside the cells, in this case just below plasma membrane. Blurred secondary electrons (SE) signal (white dashed arrows) of $TiO_2$ NCs indicates they are covered with biological layer, confirming their internalization.

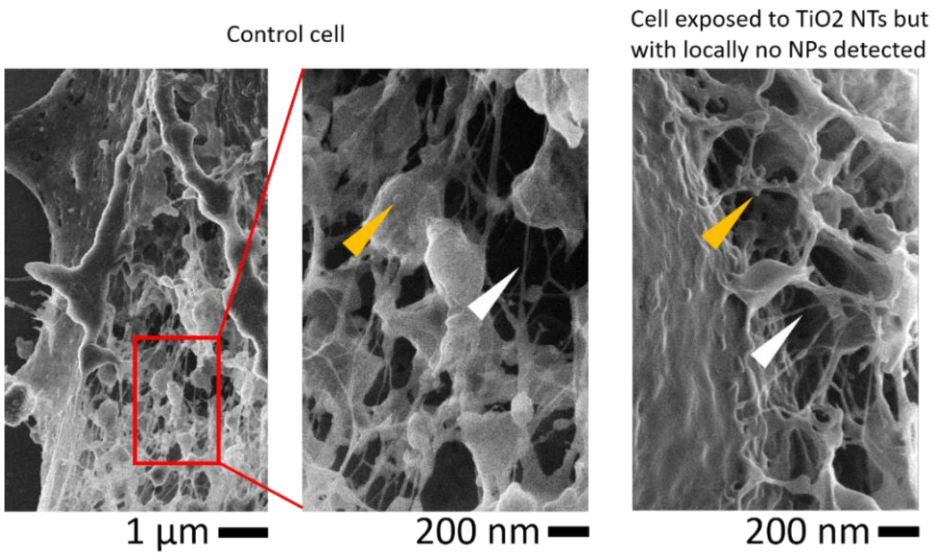


**Figure S8** Similar cellular cytoskeleton structure of the non-exposed (left) and $TiO_2$ NTs-exposed lung epithelial cells measured on the site with no chemically detected NPs (right). White arrows show the thinnest cytoskeletal structures (microfilaments) and yellow arrows the plausible ribosomes. Imaging was performed with ultra-high resolution HIM.

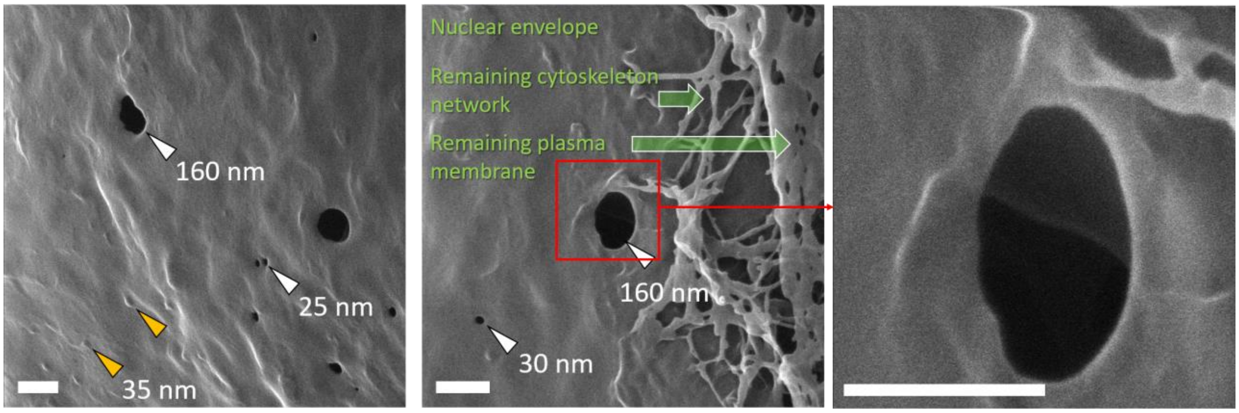


**Figure S9.** Ultra-high resolution HIM images of nuclear envelope with the density and size variability of nuclear pores (white arrows) and plausible ribosomes (yellow arrows) with remaining cytoskeleton stretching to plasma membrane (green arrow). The scale bar is 200 nm.

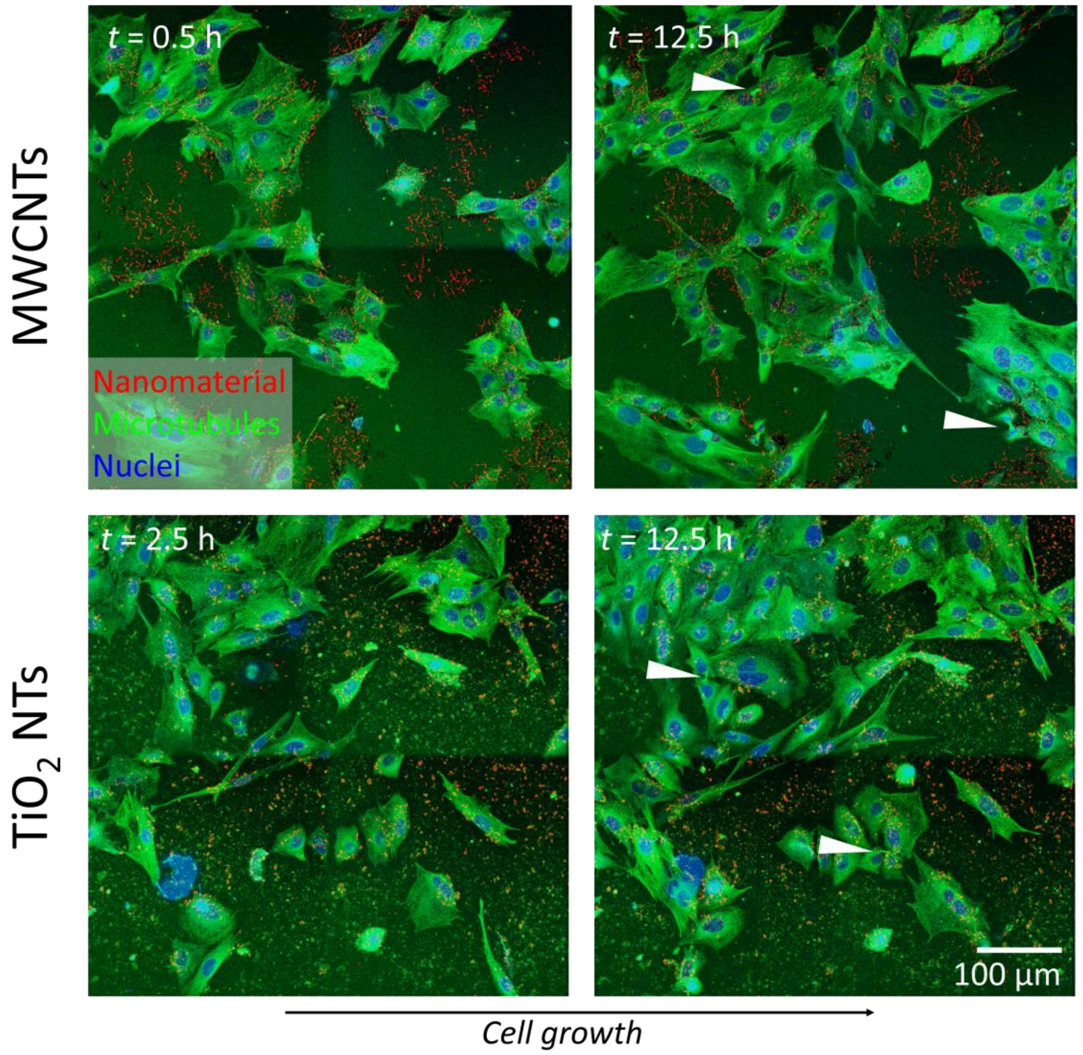


**Figure S10.** Clear cell growth and typical central spindle formation (indicated by white arrows) during the final, telophase stage of cell division shown on LA-4 cells exposed to high aspect ratio, multi-walled carbon nanotubes and $TiO_2$ nanotubes. The presented 2*2 stitched images from the time-lapse experiment were performed by a 20X magnification objective with *NA* = 0.8.

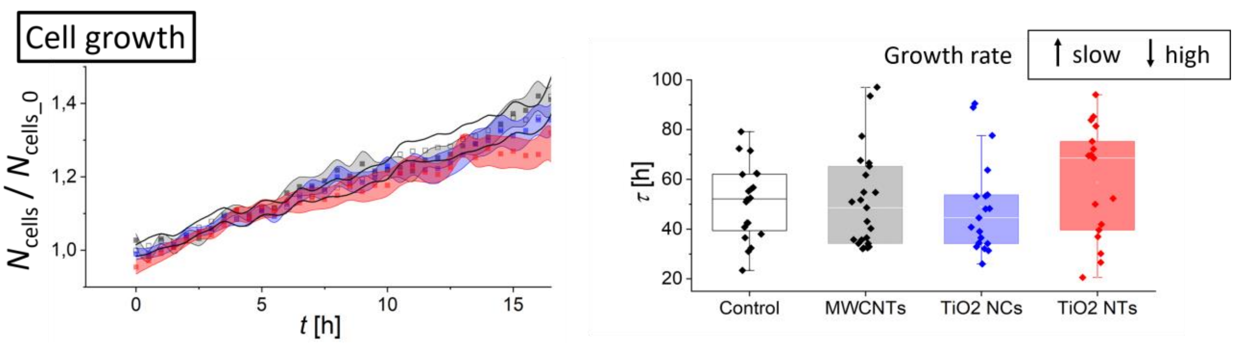

**Figure S11.** Normalized cell growth ($N_{cells}/N_{cells\_0}$) performed by an automatic measure of the number of cell nuclei throughout the first 16 h of different NPs exposure and the control with the spread representing standard error (SEm) from 24 measurements (left) and the distribution of $\tau$ fitted with the exponential curve from these 24 ROIs of 300*300μm FoV using a 20X magnification objective with $NA$ = 0.8 (right).

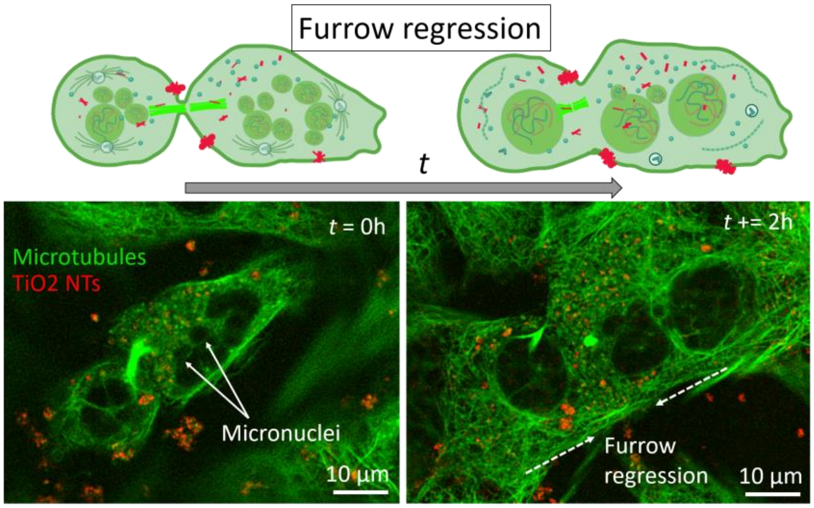


**Figure S12.** The formation of multiple micronuclei (white arrows) resulting from mitotic slippage (left) followed by a furrow regression (right) causing aneuploidy [46], after lung epithelial cells exposure to $TiO_2$ NTs. The decrease in the number of micronuclei could suggest on the nuclear envelope (NE) collapse, a key event for micronuclei dysfunction in solid tumors [47]. Numerous small $TiO_2$ NT agglomerates scattered throughout the cytoplasm of the cell are also locally found co-localized with the structurally changed microtubule network.

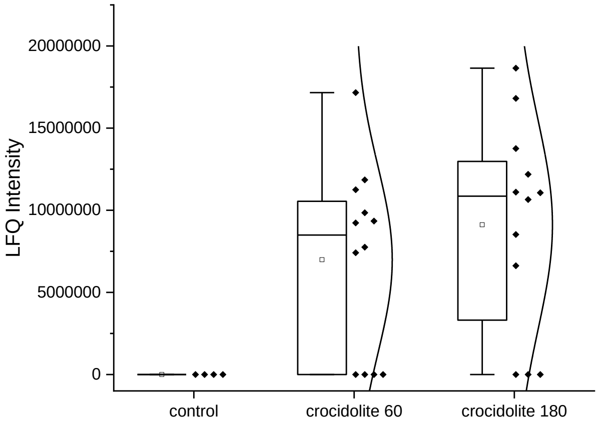

**Figure S13** Binding of septin-11 (SEPT11) found in the coronome of crocidolite (asbestos) particles 28 days after lung exposure to rats using the label-free quantification (LFQ) intensities (relative amount or the fold change compared to the control). The difference in the mean values of both concentrations of crocidolite compared to control is greater than would be expected by chance ($P$ = 0.001, Welch's t-test with equal variances not assumed).

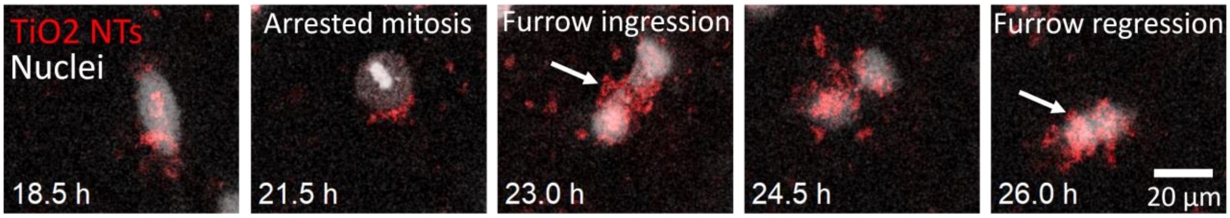


**Figure S14.** Binucleated cell formation after arrested cell mitosis and localization of TiO2 NTs within the cellular region of furrow ingression, presumably hindering the completion of cell division and ultimately resulting in furrow regression (white arrows).

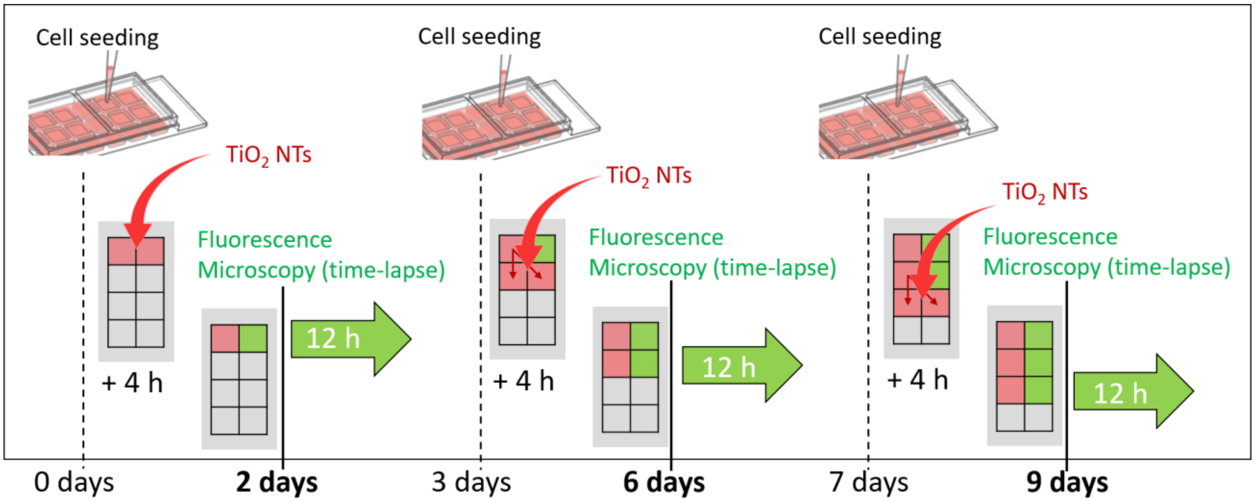


**Figure S15.** Schematics of multi-day $TiO_2$ nanotube exposure to lung epithelial LA-4 cells using surface dose ratio 10:1 coupled with a broad, few millimeter-sized field of view, time-lapse fluorescence study to examine the effects of exposure on cellular defects associated to genetic instability and carcinogenesis. At few intermediate time points of cell seeding/reseeding, suspension of $TiO_2$ NTs was added 4 hours later, once the cells had adhered sufficiently. Fluorescence microscopy was performed a few days after in time-lapse mode to easily detect cellular defects.

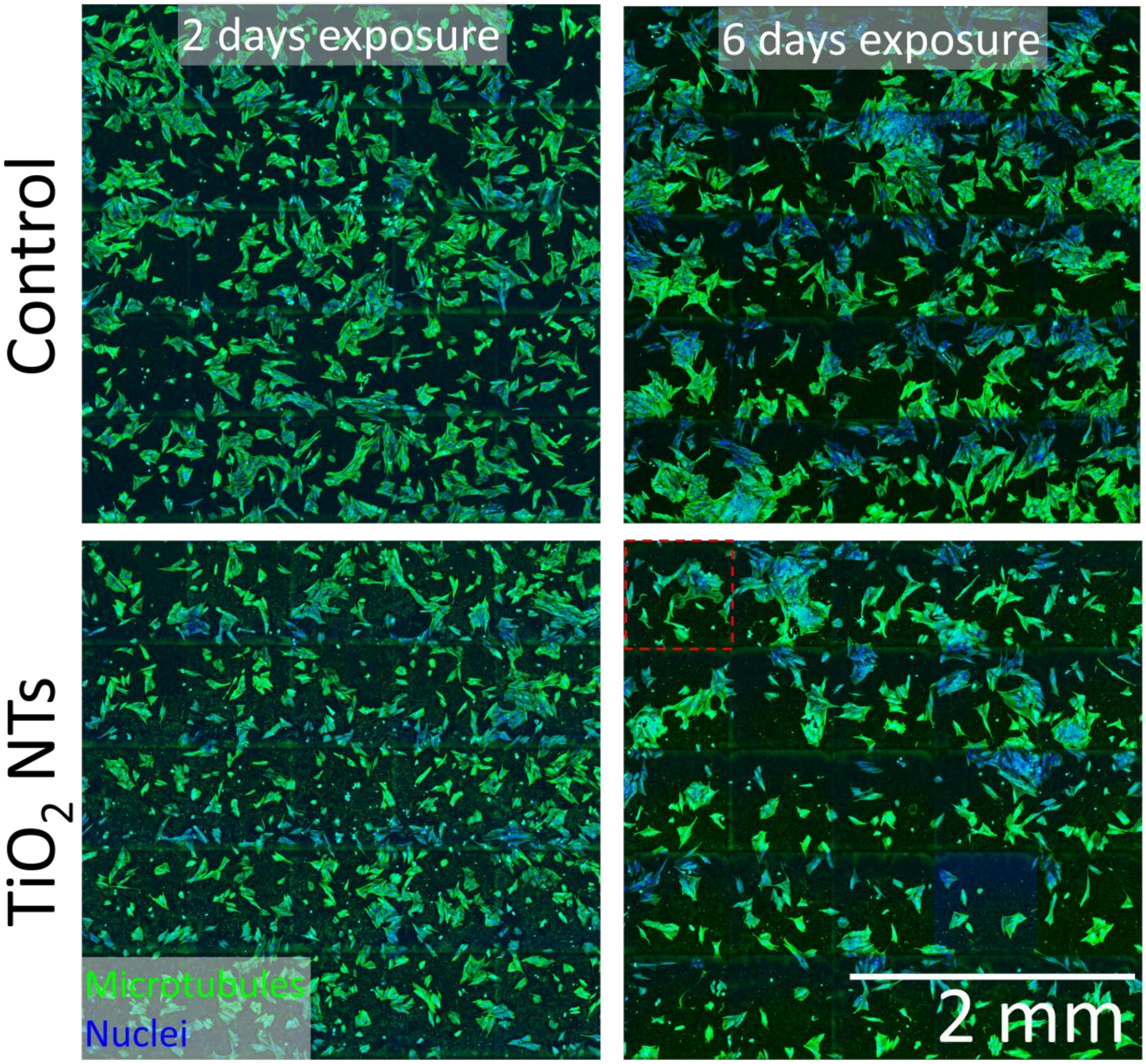


**Figure S16.** Impact of high aspect ratio $TiO_2$ nanotubes on LA-4 cell growth and cellular defects leading to cytokinesis failure measured through multi-day experiment on a broad, few $mm^2$ FoV. The presented 5*5 stitched micrographs were performed by a 10X magnification objective with $NA$ = 0.3. Horizontal stripes seen in blue are artefacts of an image stitching algorithm. An example of the automated Ilastik analysis for cell number count ($N_{cells}$) for the indicated dashed red region is shown in Figure S14.

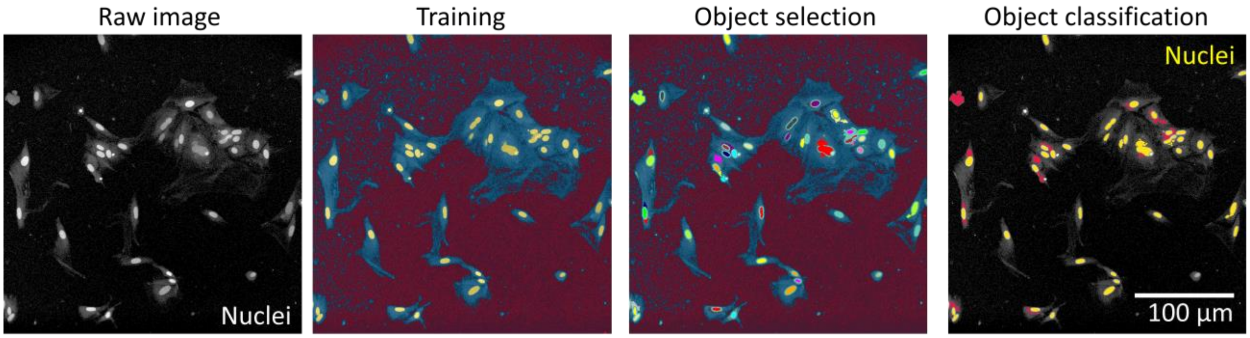


**Figure S17.** Automated Ilastik analysis of the cell number ($N_{cells}$) using appropriate training, following object selection and final object classification [21].

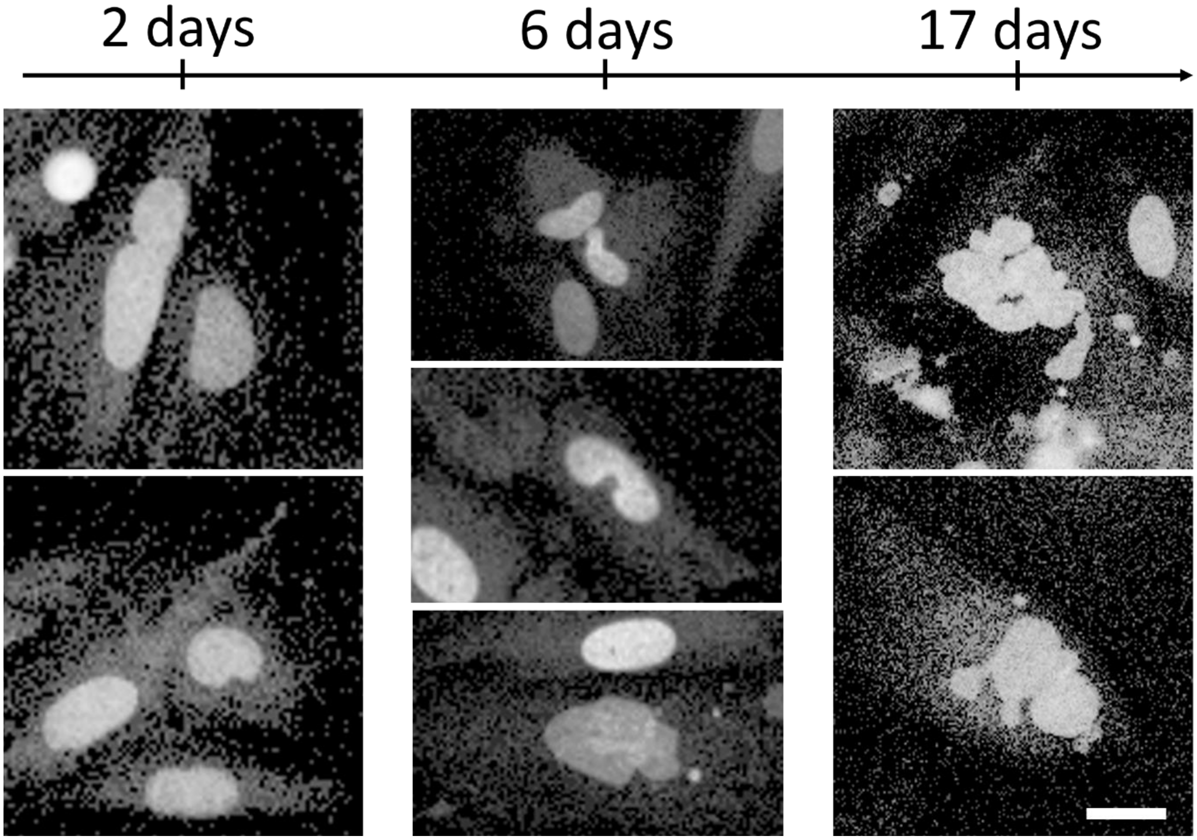


**Figure S18.** The time evolution of the formation and the extent of typical nuclear defects after $TiO_2$ nanotubes exposure to lung epithelial cells at surface dose 10:1 causing genetic instability. Nuclei were labeled with Spy DNA 555 (500 nM) prior to imaging with 10X magnification objective with *NA* = 0.3. The scale bar is 10 µm.

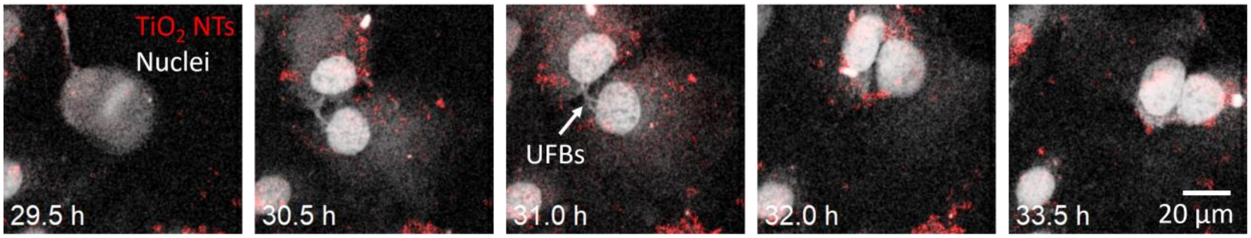


**Figure S19.** The formation of ultrafine DNA bridges (UFBs) (see white arrow) connecting the cells after metaphase stage in cell division (first time point) and prior to subsequent furrow regression, leading to bi-nuclear tetraploidy (last time point). These observations agree with the study in which furrow regression occurred exclusively in cells with chromosome bridges [48].

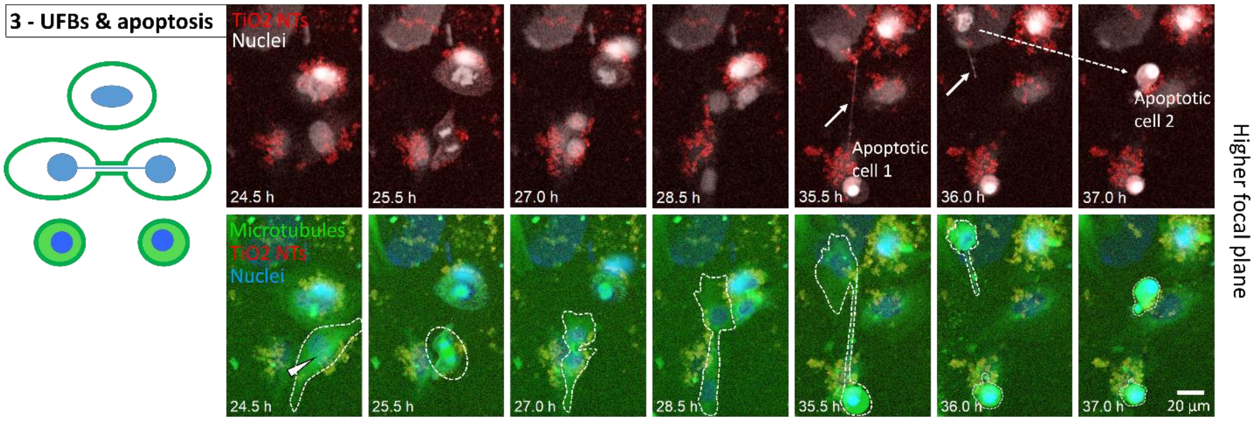


**Figure S20.** An example of cell apoptosis (dashed white arrow) following chromosomal UFBs formation (white arrow) (supporting Videos S19-20). The persistence of unresolved recombination intermediates in UFBs can induce DNA damage and chromosomal instability, and in some instances, even trigger cell apoptosis [49], similarly to our observations. The time lapse fluorescence microscopy after nuclei and microtubule labelling was performed by a 20X objective with *NA* = 0.8. $TiO_2$ NTs were detected by label-free confocal backscatter detection.

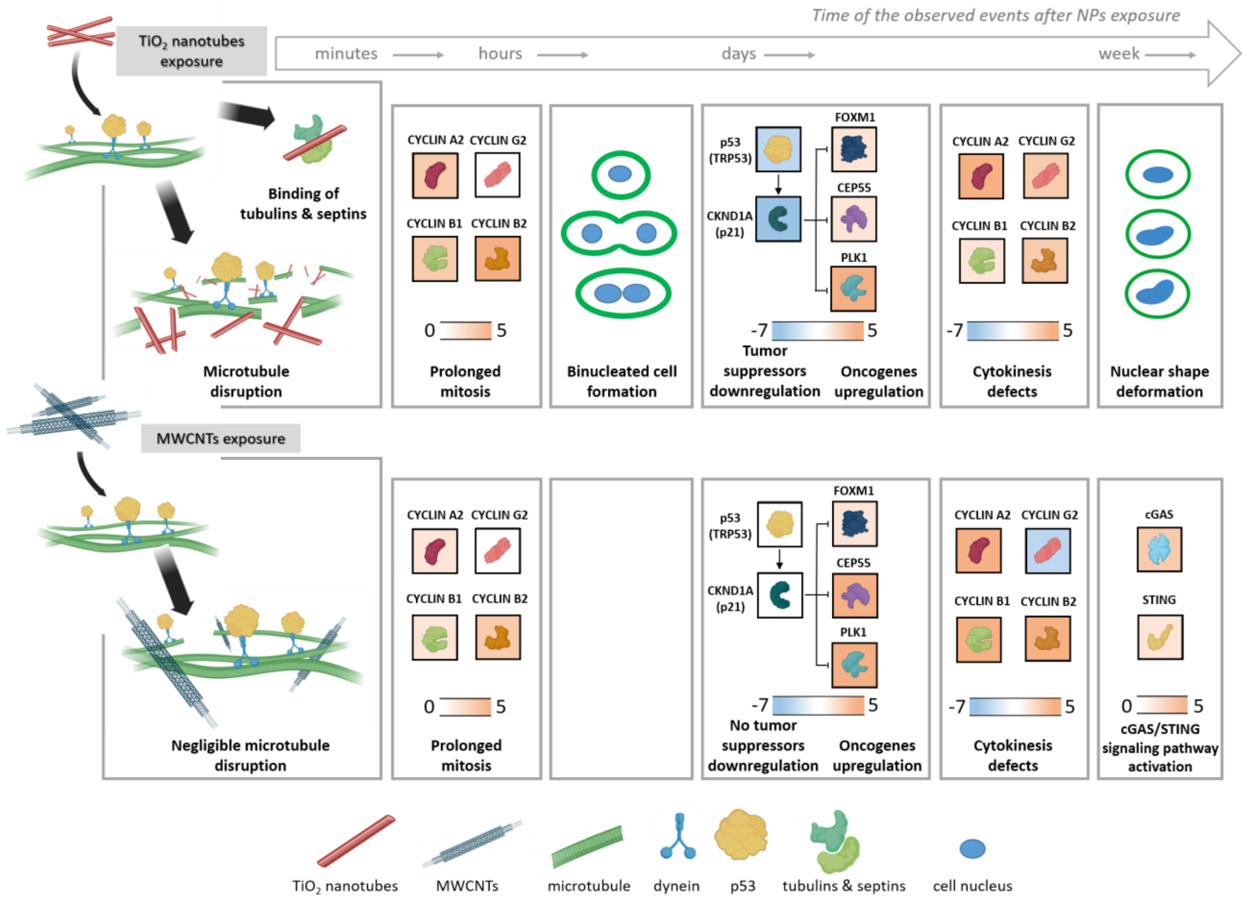


**Figure S21.** The timeline of different effects observed in lung epithelial cells in vitro that might be related to carcinogenesis of high aspect-ratio nanomaterials**.** The cells were exposed to $TiO_2$ nanotubes ($TiO_2$ NTs) and multi-walled carbon nanotubes (MWCNTs) and assessed using live-cell imaging and omics analysis. Compared to MWCNTs, $TiO_2$ NTs cause more microtubule disruption and binuclear cell formation, downregulate tumor suppressor p53, upregulate cyclin G2 2 days after exposure, and deform shape of nuclei. In contrast, MWCNTs do not influence microtubule cytoskeleton structure, downregulate cyclin G2, but upregulate the expression of tumorigenesis-related innate immune STING pathway. Abbreviations: p53 – tumor protein p53 (p53) , or transformation-related protein 53 (TRP53), this protein acts as a tumor suppressor; CKND1A (p21) – cyclin-dependent kinase inhibitor 1, alternatively p21; FOXM1 – Forkhead box protein M1, CEP55 - Centrosomal Protein 55; PLK1 – Polo-like kinase 1 (Plk1) inhibits p53 function by physical interaction and phosphorylation; STING – Stimulator of interferon genes, also known as transmembrane protein 173 (TMEM173); The color scale, represented below the expression levels of cyclins, tumour suppressors and oncogenes, depicts the log2 fold-change in mRNA expression. Graphs of non log2 transformed mRNA expression values are shown in Figure S23.

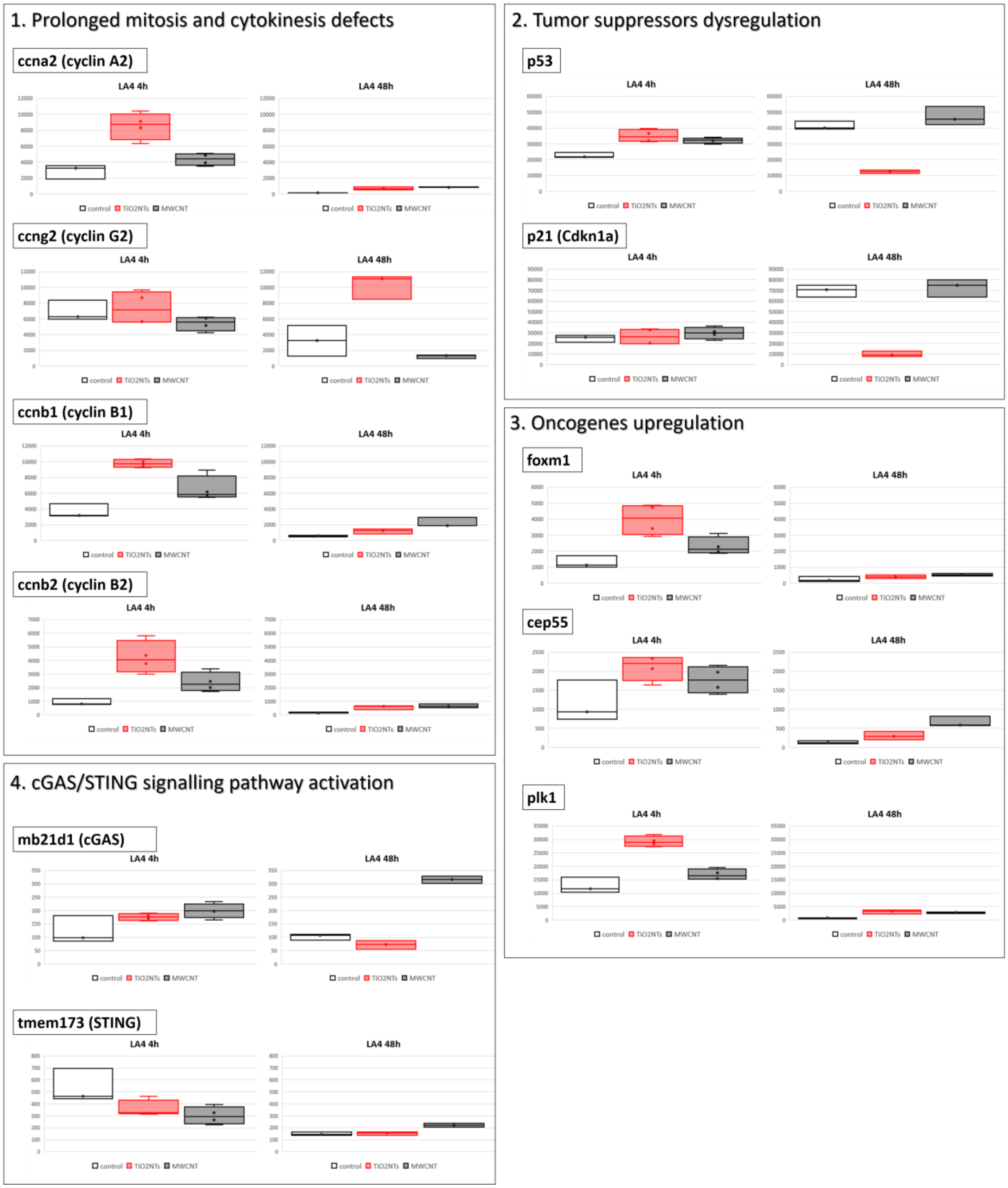


**Figure S22.** Graphs of non log2 transformed mRNA expression values reveal dysregulation of different cytokinesis and tumorigenesis related pathways after lung epithelial LA-4 cell exposure to high aspect ratio $TiO_2$ nanotubes and MWCNTs with the surface dose 10:1 for up to 2 days.

A

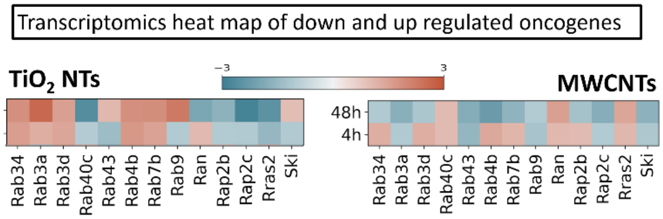


B

Oncogens found in $TiO_2$ NTs corona

| Majority protein ID | Protein names | Gene names | Ratio M | ttest |
|---|---|---|---|---|
| Q5SW88 | RAB1A, member RAS oncogene family | Rab1a | 2 | 0.06 |
| P97855 | Ras GTPase-activating protein-binding protein 1 | G3bp1 | 15207500 | 0.86 |
| Q9JKF1 | Ras GTPase-activating-like protein IQGAP1 | Iqgap1 | 3 | 0.00 |
| A2AUR7 | Ras suppressor protein 1, isoform CRA_a | Rsu1 | 8674501 | 0.08 |
| B1AWT2 | Ras-related GTP binding D, isoform CRA_a | Rragd | 6717450 | 0.03 |
| Q80X95 | Ras-related GTP-binding protein A | Rraga | 3270551 | 0.09 |
| Q91V41 | Ras-related protein Rab-14 | Rab14 | 4 | 0.79 |
| P62821 | Ras-related protein Rab-1A | Rab1A | 1600351 | 0.16 |
| P53994 | Ras-related protein Rab-2A | Rab2a | 2 | 0.34 |
| B1AQD3 | Ras-related protein Rab-34 | Rab34 | 3381801 | 0.10 |
| Q9CQD1 | Ras-related protein Rab-5A | Rab5a | 6105000 | 0.01 |
| P35278 | Ras-related protein Rab-5C | Rab5c | 2 | 0.36 |
| P51150 | Ras-related protein Rab-7a | Rab7a | 2 | 0.12 |
| P63321 | Ras-related protein Ral-A | Rala | 23044500 | 0.00 |

**Figure S23.** Transcriptomics data of up- and down-regulated distinct oncogens, particularly Rab genes known to regulate survival pathways in cancer [39] and proteomics data of oncogenesis-associated Rab proteins found in TiO2 NTs corona after 48 h exposure.
